# Protective role of neuronal and lymphoid cannabinoid CB2 receptors in neuropathic pain

**DOI:** 10.1101/2020.05.19.103580

**Authors:** David Cabañero, Angela Ramírez-López, Eva Drews, Anne Schmöle, David M. Otte, Agnieszka Wawrzczak-Bargiela, Hector Huerga Encabo, Sami Kummer, Ryszard Przewlocki, Andreas Zimmer, Rafael Maldonado

## Abstract

Cannabinoid CB2 receptor (CB2r) agonists are potential painkillers void of psychotropic effects. Peripheral immune cells, neurons and glia express CB2r, however the involvement of CB2r from these cells in neuropathic pain remains unresolved. We explored spontaneous neuropathic pain through on-demand self-administration of the selective CB2r agonist JWH133 in wild-type and knockout mice lacking CB2r in neurons, monocytes or constitutively. Operant self-administration reflected drug-taking to alleviate spontaneous pain, nociceptive and affective manifestations. While constitutive deletion of CB2r disrupted JWH133-taking behavior, this behavior was not modified in monocyte-specific CB2r knockouts and was increased in mice defective in neuronal CB2r knockouts suggestive of increased spontaneous pain. Interestingly, CB2r-positive lymphocytes infiltrated the injured nerve and possible CB2r transfer from immune cells to neurons was found. Lymphocyte CB2r depletion also exacerbated JWH133 self-administration and inhibited antinociception. This work identifies a simultaneous activity of neuronal and lymphoid CB2r that protects against spontaneous and evoked neuropathic pain.

## Introduction

Cannabinoid CB2 receptor (CB2r) agonists show efficacy in animal models of chronic inflammatory and neuropathic pain, suggesting that they may be effective inhibitors of persistent pain in humans (Bie et al., 2018; Maldonado et al., 2016; Shang and Tang, 2017). However, many preclinical studies assess reflexive-defensive reactions to evoked nociceptive stimuli and fail to take into account spontaneous pain, one of the most prevalent symptoms of chronic pain conditions in humans (Backonja and Stacey, 2004; Mogil et al., 2010; Rice et al., 2018) that triggers coping responses such as painkiller consumption. As a consequence, conclusions drawn from animal models relying on evoked nociception may not translate into efficient pharmacotherapy in humans (Huang et al., 2018; Mogil, 2009; Percie du Sert and Rice, 2014), which underlines the need to apply more sophisticated animal models with clear translational value. Operant paradigms in which animals voluntarily self-administer analgesic compounds can provide high translatability and also identify in the same experimental approach potential addictive properties of the drugs (Mogil, 2009; Mogil et al., 2010; O’Connor et al., 2011). In this line, a previous work using a CB2r agonist, AM1241, showed drug-taking behavior in nerve-injured rats and not in sham-operated animals, suggesting spontaneous pain relief and lack of abuse potential of CB2r agonists (Gutierrez et al., 2011), although the possible cell populations and mechanisms involved remain unknown. In addition, a recent multicenter study demonstrated off-target effects of this compound on anandamide reuptake, calcium channels and serotonin, histamine and kappa opioid receptors (Soethoudt et al., 2017).

CB2r, the main cannabinoid receptors in peripheral immune cells (Fernández-Ruiz et al., 2007; Schmöle et al., 2015a), are found in monocytes, macrophages and lymphocytes, and their expression increases in conditions of active inflammation (Schmöle et al., 2015b; Shang and Tang, 2017). The presence of CB2r in the nervous system was thought to be restricted to microglia and limited to pathological conditions or intense neuronal activity (Manzanares et al., 2018). However, recent studies using electrophysiological approaches and tissue-specific genetic deletion revealed functional CB2r also in neurons, where they modulate dopamine-related behaviors (Zhang et al., 2014) and basic neurotransmission (Quraishi and Paladini, 2016; Stempel et al., 2016). Remarkably, the specific contribution of immune and neuronal CB2r to the development of chronic pathological pain has not yet been established.

This work investigates the participation of neuronal and non-neuronal cell populations expressing CB2r in the development and control of chronic neuropathic pain. We used a pharmacogenetic strategy combining tissue-specific CB2r deletion and drug self-administration to investigate spontaneous neuropathic pain. Constitutive and conditional knockouts lacking CB2r in neurons or monocytes were nerve-injured, subjected to operant self-administration of the specific CB2r agonist JWH133 (Soethoudt et al., 2017) and were evaluated for nociceptive and anxiety-like behavior. We also explored infiltration of CB2r-positive immune cells in the injured nerve of mice receiving bone marrow transplants from CB2-GFP BAC mice. Finally, immunological blockade of lymphocyte extravasation was used to investigate the effect of this cell type on spontaneous neuropathic pain and its involvement on the pain-relieving effects of the cannabinoid CB2r agonist.

## Results

### Self-administration of a CB2r agonist to alleviate spontaneous pain and anxiety associated behavior

CB2r agonists have shown efficacy reducing evoked sensitivity and responses of negative affect in mouse models of chronic pain (Maldonado et al., 2016). Although antinociception is a desirable characteristic for drugs targeting chronic neuropathic pain, it is unclear whether the pain-relieving effects of the CB2r agonist would be sufficient to elicit drug-taking behavior in mice and the cell populations involved. To answer these questions, mice underwent a PSNL or a sham surgery and were placed in operant chambers where they had to nose poke on an active sensor to obtain i.v. self-administration of the CB2r agonist JWH133 or vehicle (Figure 1A). Sham mice or nerve-injured animals receiving vehicle or the low dose of JWH133 (0.15 mg/kg/inf) did not show significant differences in active nose-poking during the last 3 days of the drug self-administration period (Figure 1B, Figure 1-figure supplement 1A). Conversely, nerve-injured mice exposed to the high dose of JWH133 (0.3 mg/kg/inf) showed higher active responses than sham-operated mice receiving the same treatment (Figure 1B, Figure 1-figure supplement 1A). As expected, the operant behavior of sham-operated mice exposed to JWH133 was not different from that of sham mice exposed to vehicle, suggesting absence of reinforcing effects of the CB2r agonist in mice without pain (Figure 1B, Figure 1-figure supplement 1A). The number of nose pokes on the inactive sensor was similar among the groups, indicating absence of locomotor effects of the surgery or the pharmacological treatments. Thus, operant JWH133 self-administration was selectively associated to the neuropathic condition.

**Figure 1.**
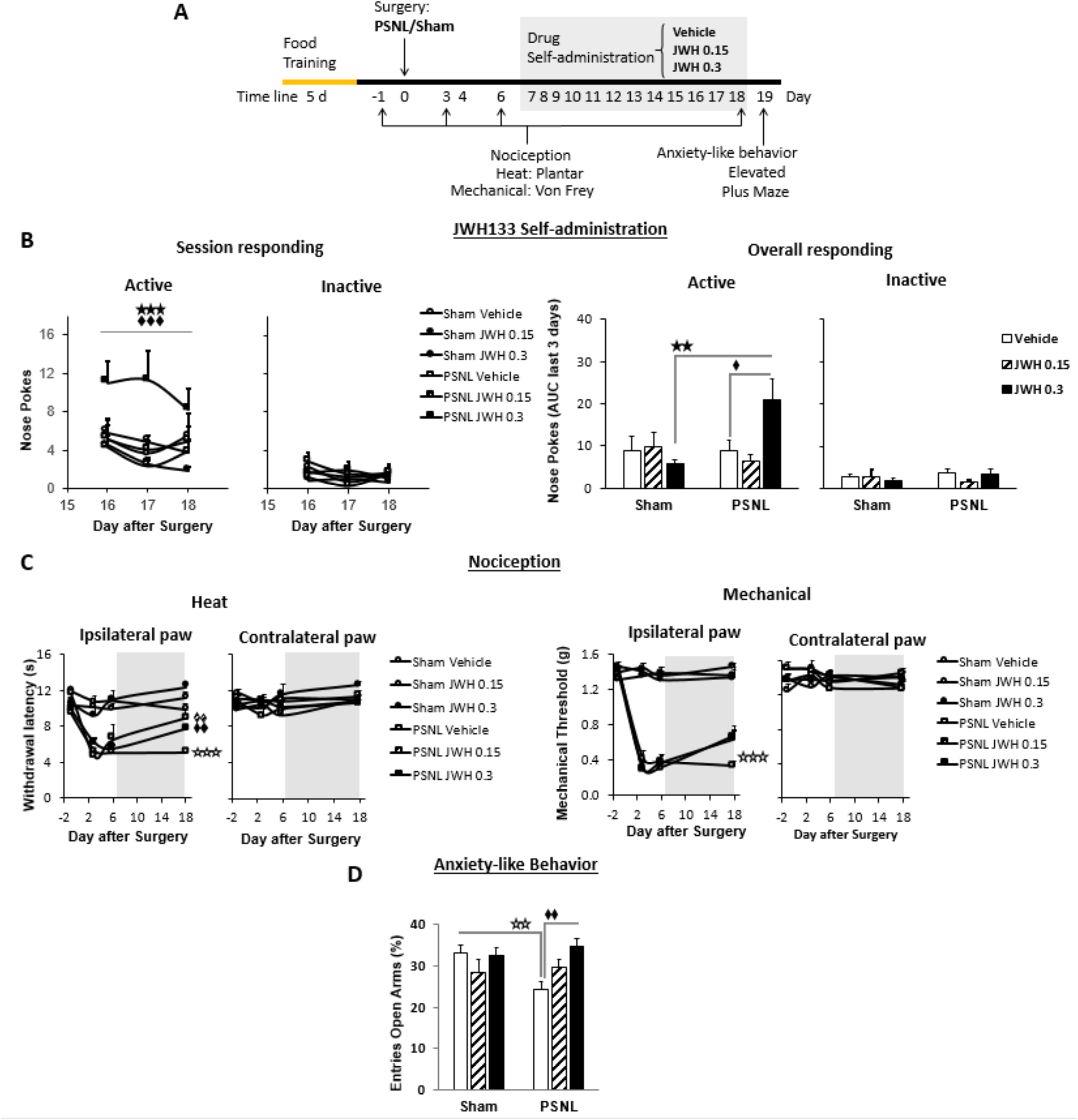
C57BL/6J mice self-administer a CB2r agonist with antinociceptive and anxiolytic-like properties. **A)** Timeline of the drug self-administration paradigm. Mice were trained in Skinner boxes (5 days, 5d) where nose-poking an active sensor elicited delivery of food pellets. Partial sciatic nerve ligation (PSNL) or sham surgery were conducted (day 0) followed by jugular catheterization to allow intravenous (i.v.) drug infusion. From days 7 to 18, mice returned to the operant chambers and food was substituted by i.v. infusions of JWH133 (0.15 or 0.3 mg/kg/inf.). Mechanical and thermal sensitivity were assessed before (−1) and 3, 6 and 18 days after PSNL using Plantar and von Frey tests. Anxiety-like behaviour was measured at the end (day 19) with the elevated plus maze. **B)** Nerve-injured mice poked the active sensor to consume the high dose of JWH133 (0.3 mg/kg/inf.). **C)** PSNL induced ipsilateral thermal and mechanical sensitization (days 3 and 6). JWH133 inhibited thermal hypersensitivity but the effect on mechanical nociception was not significant **D)** Nerve-injured mice receiving vehicle showed decreased percentage of entries to the open arms of the elevated plus maze, whereas PSNL mice receiving JWH133 0.3 mg/kg/inf. did not show this alteration. N=5-10 mice per group. Shaded areas represent drug self-administration. Mean and error bars representing SEM are shown. Stars represent comparisons vs. sham; diamonds vs. vehicle. *p<0.05; **p<0.01; ***p<0.001. Figure 1–figure supplement 1

Nociceptive responses to thermal and mechanical stimuli were assessed before and after the self-administration period (days -1, 3, 6 and 18). Before the treatment with the CB2r agonist, all nerve-injured mice developed heat and mechanical hypersensitivity in the ipsilateral paw (Figure 1C). After self-administration (shaded area, Figure 1C) mice exposed to JWH133 showed a significant reduction in heat hypersensitivity (Figure 1C, day 18, ipsilateral paw), although the alleviation of mechanical hypersensitivity did not reach statistical significance in this experiment. No significant drug effects were observed in the contralateral paws.

We also studied affective-like behavior in mice exposed to this chronic pain condition. Anxiety-like behavior was enhanced in nerve-injured mice treated with vehicle, as these mice visited less frequently the open arms of the elevated plus maze (Figure 1D). This emotional response was absent in nerve-injured mice repeatedly exposed to the high dose of JWH133 (Figure 1D). Therefore, the high dose of JWH133 elicited a drug-taking behavior selectively associated to spontaneous pain relief, and had efficacy limiting the pronociceptive effects of the nerve injury and its emotional-like consequences.

### CB2 receptor mediates JWH133 effects on spontaneous pain alleviation

JWH133 has been recently recommended as a selective CB2r agonist to study the role of CB2r in biological and disease processes due to its high selectivity for this receptor (Soethoudt et al., 2017). To investigate the specificity of the CB2r agonist in our model, the high dose of JWH133 (0.3 mg/kg/inf) was offered to nerve-injured mice constitutively lacking the CB2r (CB2KO) and to C57BL/6J wild-type mice. CB2KO mice showed a significant disruption of JWH133-taking behavior on the last sessions of the drug self-administration period (Figure 2A, Figure 2-figure supplement 1A). Overall discrimination between the active and inactive sensors was also significantly blunted in CB2KO mice (Source Data File) and inactive nose pokes were similar in both groups of mice, indicating absence of genotype effect on locomotion (Figure 2A, Figure 2-figure supplement 1A). The disruption of drug-taking behavior shown in CB2KO mice was accompanied by an inhibition of JWH133 effects on nociceptive and affective behavior (Figure 2B, Figure 2C).

**Figure 2.**
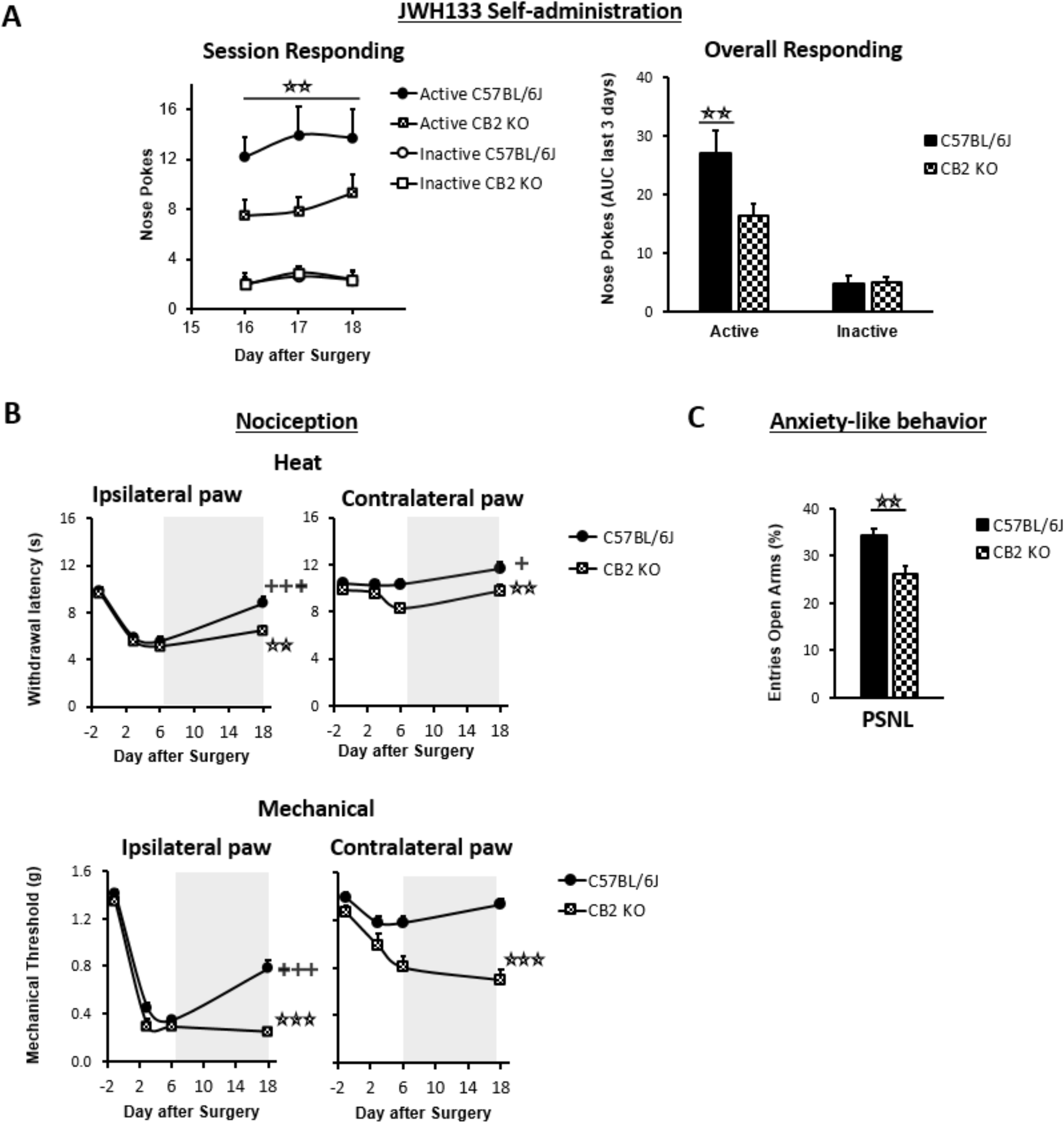
Nerve-injured mice constitutively lacking CB2r show disruption of JWH133 intake and blunted effects of the drug. CB2r constitutive knockout mice (CB2 KO) and C57BL/6J mice were food-trained in Skinner boxes (Food training, 5 days), subjected to a partial sciatic nerve ligation (PSNL, day 0), catheterized and exposed to high doses of the CB2r agonist JWH133 (0.3 mg/kg/inf., days 7 to 18). Nociceptive sensitivity to heat (Plantar) and mechanical (von Frey) stimulation were measured before and after the nerve injury (−1,3,6,18), and anxiety-like behaviour was evaluated at the end (day 19). **A)** CB2 KO mice showed decreased active operant responding for the CB2r agonist. **B)** The effects of JWH133 on thermal nociception were reduced in constitutive knockout mice. CB2 KO mice showed contralateral mechanical and thermal sensitization and complete abolition of JWH133 effects on mechanical hypersensitivity. **C)** Anxiety-like behaviour after the treatment worsened in CB2 KO mice. N=16-19 mice per group. Mean and error bars representing SEM are shown. Shaded areas represent drug self-administration. Stars represent comparisons vs. C57BL/6J mice; crosses represent day effect. *p<0.05; **p<0.01; ***p<0.001. Figure 2–figure supplement 1

CB2KO and C57BL/6J mice developed similar thermal and mechanical hypersensitivity in the injured paw (Figure 2B, day 6, Ipsilateral paw), although CB2KO mice also developed hypersensitivity in the contralateral paw, as previously described (Racz et al., 2008). While C57BL/6J mice showed significant recovery of thermal and mechanical thresholds after JWH133 self-administration (Figure 2B, day 18), CB2KO mice showed no effects of the treatment on mechanical sensitivity (Figure 2B, day 18, Mechanical) and a partial recovery of the thresholds to heat stimulation (Figure 2B, day 18, Heat). Contralateral mechanical sensitization was still present in CB2KO mice exposed to the CB2r agonist (Figure 2B, Contralateral paw). Likewise, nerve-injured C57BL/6J mice showed less anxiety-like behavior after JWH133 self-administration than CB2KO mice (Figure 2C), suggesting that these anxiolytic-like effects of JWH133 are mediated by CB2r. Hence, CB2KO mice showed reduced drug-taking behavior accompanied by blunted inhibition of JWH133 effects on mechanical nociception and anxiety-like behavior, confirming mediation of these effects by CB2r.

### Participation of neuronal and monocyte CB2r in neuropathic pain symptomatology

CB2r were initially described in peripheral immune cells (Munro et al., 1993), although they have been found in multiple tissues including the nervous system. In order to distinguish the participation of CB2r from different cell types on spontaneous neuropathic pain, we conducted the self-administration paradigm in nerve-injured mice lacking CB2r in neurons (Syn Cre+ mice) or in monocyte-derived cells (LysM Cre+) and in their wild-type littermates (Cre Neg). Syn Cre+ mice showed increased active operant responding for JWH133 (Figure 3A, Figure 3-figure supplement 1A), suggesting increased spontaneous pain and possible decrease of drug effects. On the other hand, LysM Cre+ mice did not show significant alteration of drug-taking behavior (Figure 3A, Figure 3-figure supplement 1A). Inactive responding was also similar between Cre Neg and knockout mice. Thus, data from the drug self-administration experiments showed persistence of drug effects in the different genotypes and increased self-administration in mice lacking neuronal CB2r, suggestive of increased spontaneous pain.

**Figure 3.**
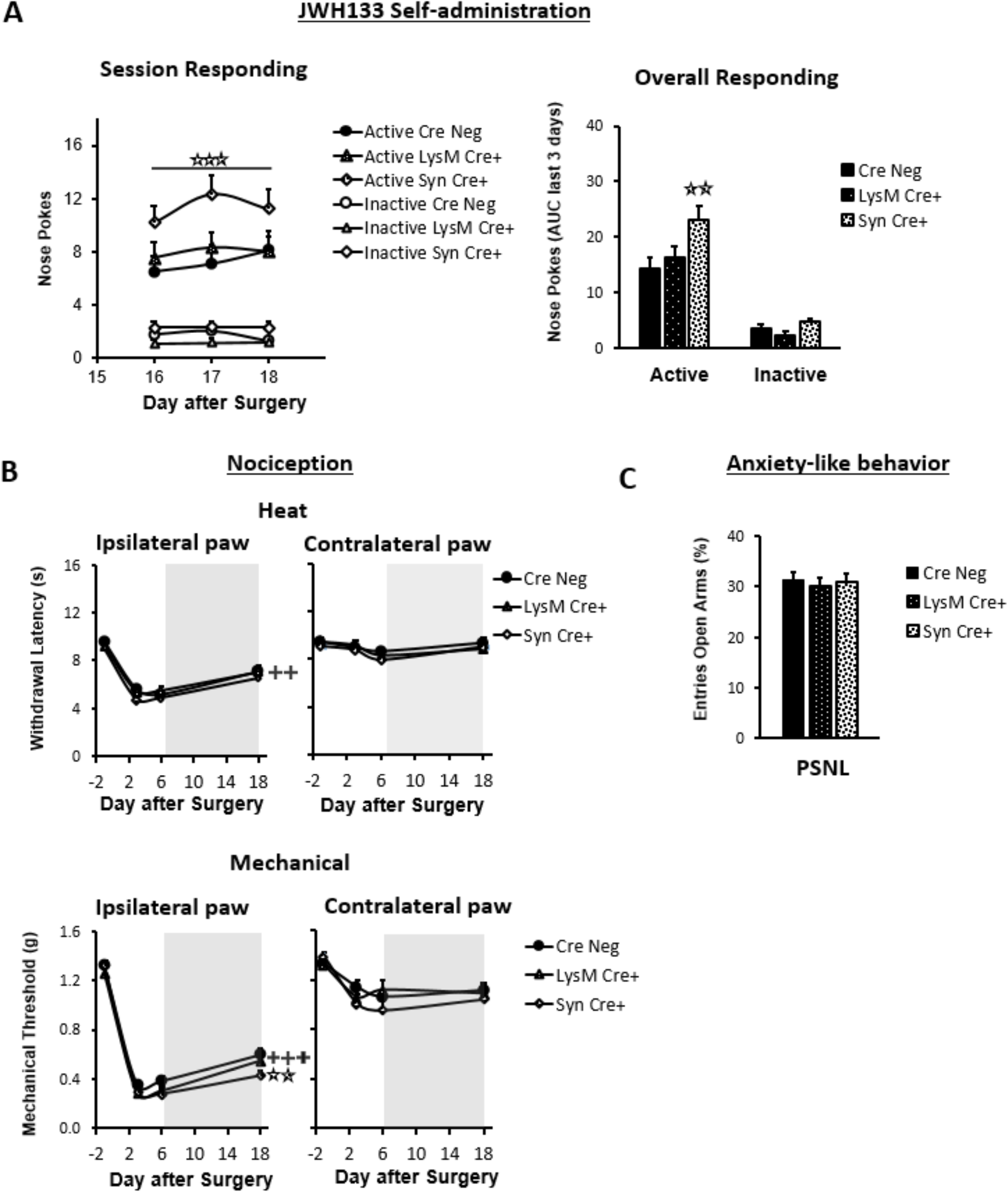
Nerve-injured mice defective in neuronal CB2r show increased self-administration of the CB2r agonist JWH133 and a decrease in the antinociceptive effects of the drug. Mice lacking CB2r in neurons (Syn Cre+), in monocytes (LysM Cre+) or their wild-type littermates (Cre Neg) were food-trained in Skinner boxes (Food training, 5 days), subjected to partial sciatic nerve ligation (PSNL, day 0), catheterized and exposed to JWH133 (0.3 mg/kg/inf., days 7 to 18). Nociceptive sensitivity to heat (Plantar) and mechanical (von Frey) stimulation were measured before and after nerve injury (−1,3,6,18), anxiety-like behaviour was evaluated at the end (day 19). **A)** Syn Cre+ mice showed increased active operant responding for JWH133 in the last sessions of the self-administration period **B)** All mouse strains showed decreased heat nociception after JWH133 treatment, and Syn Cre+ mice showed reduced effects of JWH133 on mechanical nociception. **C)** Every mouse strain showed similar anxiety-like behavior after JWH133 self-administration. No significant differences were found between LysM Cre+ and Cre Neg mice. N=18-36 mice per group. Mean and error bars representing SEM are shown. Shaded areas represent drug self-administration. Stars represent comparisons vs. Cre Neg mice; crosses represent day effect. *p<0.05; **p<0.01; ***p<0.001. Figure 3–figure supplement 1

We also measured antinociceptive and anxiolytic-like effects of JWH133 self-administration (Figure 3B, Figure 3C). The three mouse lines showed similar evoked responses to nociceptive stimulation after nerve injury (Figure 3B). A slight but significant impairment on the effect of JWH133 on mechanical sensitivity was found in Syn Cre+ mice (Figure 3C) in spite of the increased JWH133 consumption, compatible with reduced efficacy of JWH133 in this mouse strain. The assessment of anxiety-like behavior did not reveal apparent differences among the three genotypes (Figure 3C). Thus, the increased JWH133 consumption observed in Syn Cre+ mice was not reflected in increased anxiolysis and JWH133 antinociceptive effects were blunted, suggesting partial involvement of neuronal CB2r in the development of spontaneous and evoked neuropathic pain.

### Infiltration of non-neuronal CB2r-GFP+ cells in the injured nerve

The persistence of JWH133 effects after genetic deletion of CB2r from neurons and monocyte-derived cells led us to hypothesize that CB2r of other cell types may still exert neuromodulatory effects. To investigate possible infiltration of non-neuronal GFP+ cells in the injured nerve, we transplanted bone marrow cells from C57BL/6J or CB2r-GFP BAC mice to lethally irradiated CB57BL/6J recipient mice (Figure 4-figure supplement 1). Mice transplanted with bone marrow from CB2r-GFP mice (CB2r-GFP BMT) or from C57BL/6J mice (C57BL/6J BMT) were exposed to a partial sciatic nerve ligation or a sham surgery and dorsal root ganglia were collected 14 days later. A significant infiltration of non-neuronal GFP+ cells was revealed in nerve injured CB2r-GFP BMT mice (∼34 cells/mm^2^, Figure 4A, Figure 4-figure supplement 2), indicating that CB2r-expressing cells invaded the injured nerve. Immunostaining to identify these cell types revealed co-localization with macrophage and lymphocyte markers. Nearly 60% of infiltrating macrophages and around 40% of the lymphocytes were found to be GFP+ (Figure 4B, Figure 4C, Figure 4-figure supplement 3). Surprisingly, a significant percentage of neurons was also found to express GFP in CB2r-GFP BMT mice (Figure 4D). The percentage of GFP+ neurons was higher in nerve-injured mice (∼4% of total neurons) than in sham-operated animals (∼2%, Figure 4D, Figure 4-figure supplement 4). Since GFP could only come from bone-marrow transplanted cells, this finding suggests a transfer of CB2r from bone-marrow derived cells to neurons. Hence, nerve injury facilitated the invasion of affected ganglia by CB2r-positive immune cells and promoted a neuronal GFP expression compatible with transfer of CB2r from immune cells to neurons.

**Figure 4.**
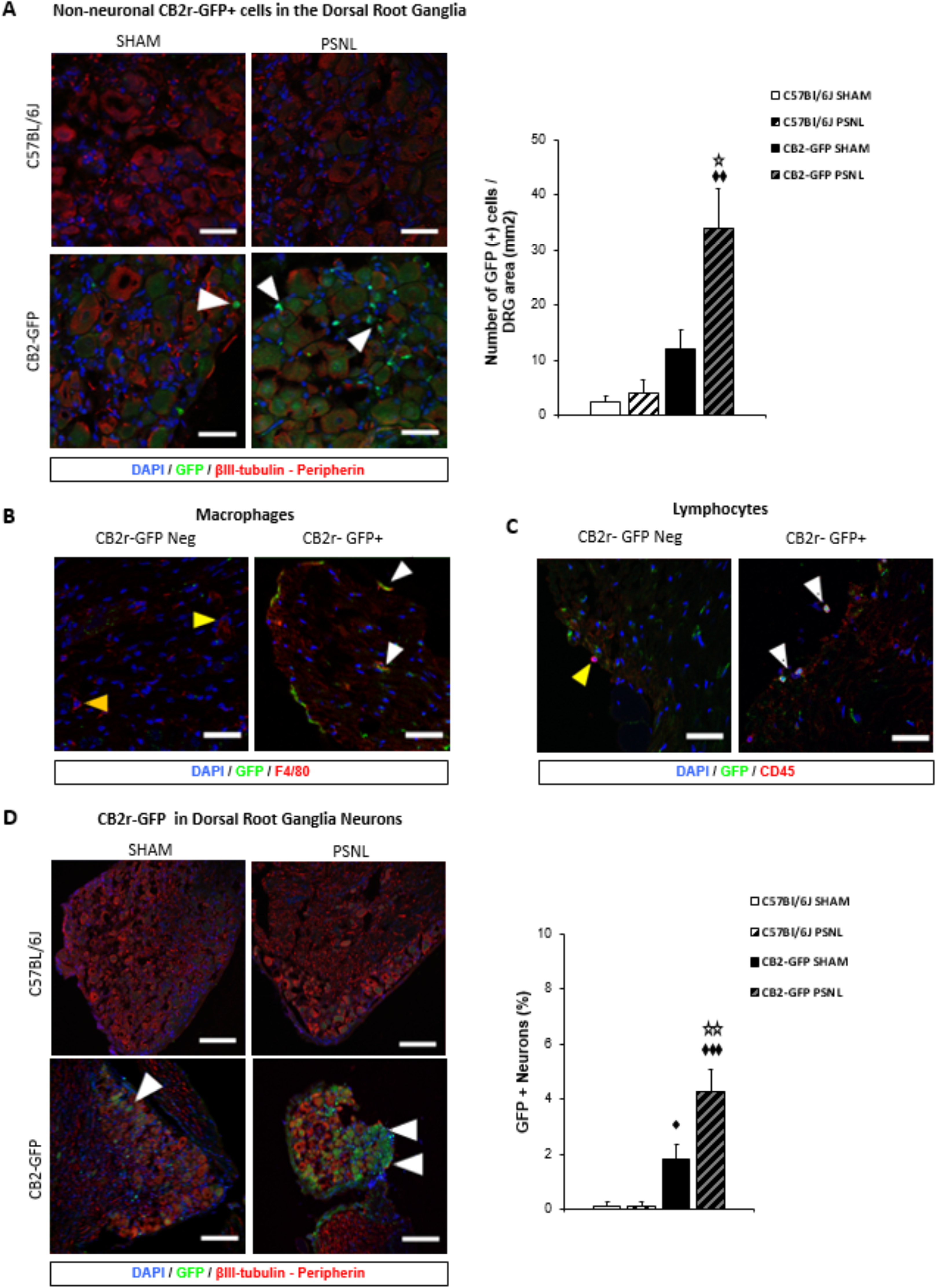
CB2r-GFP immune cells infiltrate the dorsal root ganglia of the injured nerve and GFP from bone-marrow derived cells is also found inside peripheral neurons. The figure shows images of L3-L5 dorsal root ganglia from sham (SHAM) or nerve-injured mice (PSNL) transplanted with bone marrow cells from CB2 GFP BAC mice (CB2-GFP) or C57BL6/J mice (C57BL6/J). **A, D)** Dorsal root ganglia sections stained with the nuclear marker DAPI, anti-GFP, and neuronal markers anti-β-III tubulin and anti-peripherin. **A)** CB2-GFP mice showed significant infiltration of GFP+ bone marrow-derived cells after the nerve injury, whereas sham or nerve-injured C57BL6/J mice did not show significant GFP immunorreactivity. Split channels in Figure 4-figure supplement 2. **B)** Co-localization of CB2-GFP and the macrophage marker anti-F4/80. Co-staining with anti-GFP and anti-F4/80 revealed GFP+ (∼60%) and GFP negative macrophages infiltrating the injured nerve. Split channels in Figure 4-figure supplement 3A. **C)** Co-staining with anti-GFP and anti-CD45 revealed GFP+ (∼40%) and GFP negative lymphocytes infiltrating the injured nerve. Split channels in Figure 4-figure supplement 3B. **D)** CB2-GFP mice showed a percentage of GFP+ neurons that was enhanced with the nerve injury. Scale bar, 140 μm. Split channels in Figure 4-figure supplement 4. Scale bar for B), C), D), 45 μm. Yellow arrows point to GFP negative cells and white arrows to GFP+ cells. A certain degree of image processing has been applied equally across the entire merged images for optimal visualization. N=2-3 mice per group. Means and error bars representing SEM are shown. Stars represent comparisons vs. sham; diamonds vs. C57BL6/J. *p<0.05, **p<0.01, ***p<0.001. Figure 4–figure supplement 1 Figure 4–figure supplement 2 Figure 4–figure supplement 3 Figure 4–figure supplement 4

### Lymphocyte involvement on JWH133 efficacy

The discovery of CB2r-expressing lymphocytes invading the dorsal root ganglia of nerve-injured mice prompted us to investigate the role of this cell type in spontaneous neuropathic pain. To answer this question C57BL/6J mice were repeatedly treated with a control IgG or with an antibody targeting intercellular adhesion molecule 1 (ICAM1), a protein required for lymphocyte extravasation (Labuz et al., 2009). Mice under treatment with anti-ICAM-1 or with the control IgG were exposed to JWH133 self-administration. Instead of reducing the intake of the CB2r agonist, anti-ICAM1 significantly increased active nose poking to obtain i.v. JWH133 without altering the inactive nose poking (Figure 5A, Figure 5-figure supplement 1A), suggesting increased spontaneous pain. This result is in agreement with previous works showing protection against chronic inflammatory and neuropathic pain mediated by lymphoid cells (Labuz et al., J Clin Invest 2009; Baddack-Werncke et al., J Neuroinflammation 2017). Interestingly, thermal and mechanical nociception before self-administration were similar in anti-ICAM1 and control IgG-treated mice (Figure 5B). After self-administration, the alleviation of thermal sensitivity was similar in control IgG and anti-ICAM1-treated mice (Figure 5B), but mice treated with anti-ICAM1 also showed an abolition of the antinociceptive effect of JWH133 on mechanical sensitivity (Figure 5B). This was evident in spite of the increased drug-taking behavior shown by mice treated with anti-ICAM1 (Figure 5A), which reveals decreased antinociceptive efficacy of JWH133 in these mice. On the contrary, anxiety-like behavior was similar in Control IgG and anti-ICAM1 mice (Figure 5C). To confirm an effect of the antibody treatment on lymphocyte infiltration, RT-PCR for white blood cell markers was performed in the dorsal root ganglia of mice subjected to the behavioral paradigm. As expected, a significant decrease in T cell markers CD2 and CD4 was observed in mice treated with anti ICAM-1 (Figure 5D, T cell panel). Interestingly, anti ICAM-1 also showed a pronounced increase in B cell marker CD19 (Figure 5D) and no alteration of the macrophage marker C1q (Figure 5D). Hence, our results reveal that lymphoid cells are involved in spontaneous neuropatic pain and are also necessary for the antinociceptive effect of JWH133 on mechanical sensitivity.

**Figure 5.**
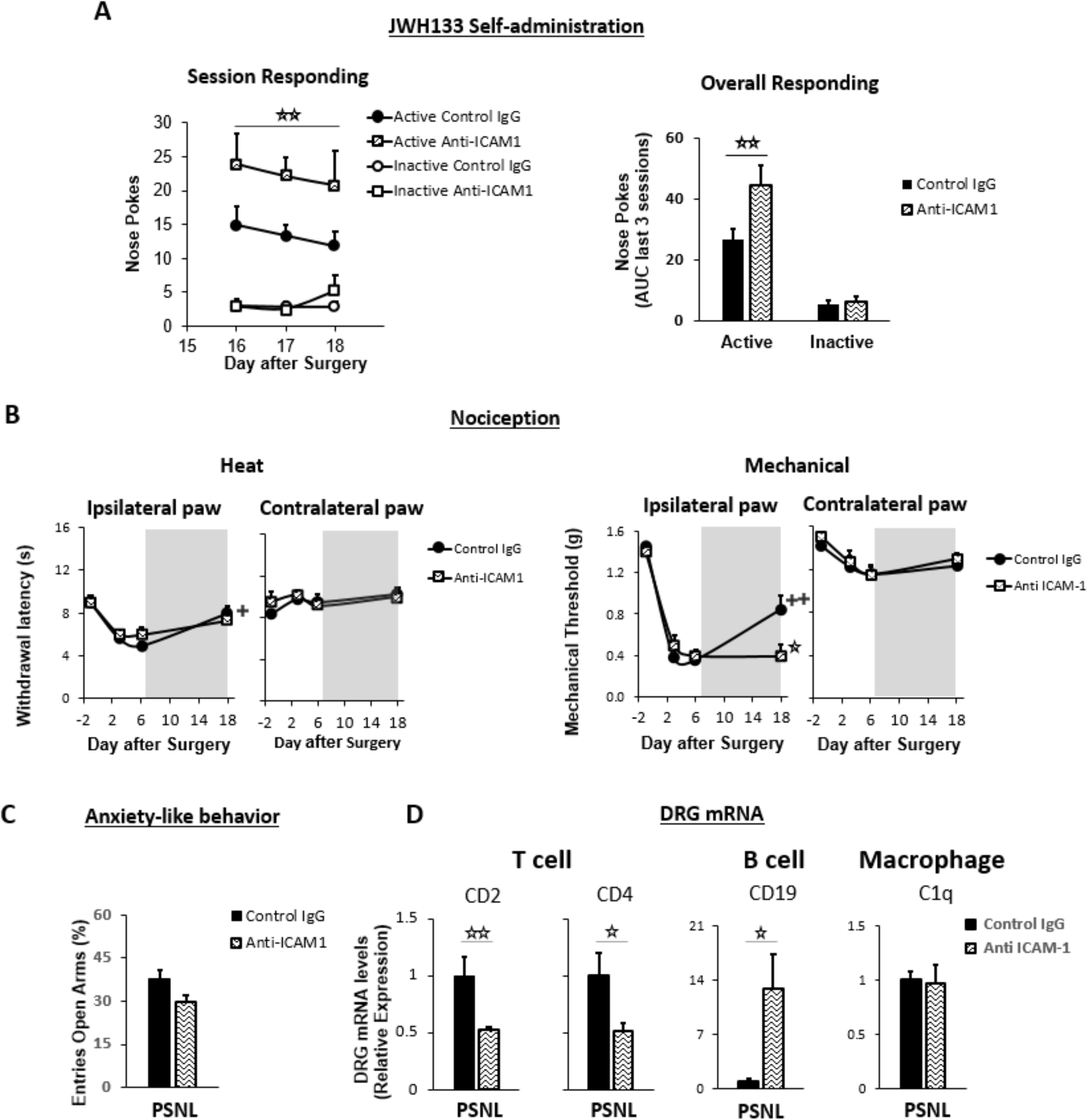
Lymphocytes modulate the effects of JWH133 on spontaneous pain and mechanical nociception. C57BL/6J mice were food-trained in Skinner boxes (Food training, 5 days), subjected to partial sciatic nerve ligation (PSNL, day 0), catheterized and exposed to high doses of the CB2r agonist JWH133 (0.3 mg/kg/inf., days 7 to 18). Treatments with Anti-ICAM1 (an antibody that inhibits lymphocyte extravasation) or control IgG were given intraperitoneally once a day from day 0 until the end of self-administration. Nociceptive sensitivity to heat (Plantar) and mechanical (von Frey) stimulation was measured before and after nerve injury (−1,3,6,18), and anxiety-like behaviour was evaluated at the end (day 19). Dorsal root ganglia were collected for mRNA analysis **A)** Mice treated with anti-ICAM1 showed increased active responding for JWH133. **B)** Thermal nociception after JWH133 self-administration was similar in mice treated with anti-ICAM1 or control IgG. Conversely, JWH133 effects on mechanical nociception were abolished by anti-ICAM1.**C)** Anxiety-like behaviour was similar in anti-ICAM1 and control IgG mice. **D)** Levels of mRNA from T cell markers CD2 and CD4 were decreased in the dorsal root ganglia of anti-ICAM1 mice. Conversely, levels of B cell marker CD19 increased. Macrophage marker C1q was unaffected. N=6-7 mice per group. Shaded areas represent drug self-administration. Mean and error bars representing SEM are shown. Stars represent comparisons vs. control IgG group; crosses indicate day effect. *p<0.05; **p<0.01; ***p<0.001. Figure 5–figure supplement 1

## Discussion

This work shows a protective function of CB2r from neurons and lymphocytes on spontaneous neuropathic pain and the involvement of these cell populations in CB2-induced antinociception, as revealed by increased self-administration of the CB2r agonist JWH133 in mice defective in lymphocyte and neuronal CB2r. Previous works already demonstrated antinociceptive and emotional-like effects of CB2r agonists in rodent models of acute and chronic pain (Gutierrez et al., 2011; Ibrahim et al., 2003; Jafari et al., 2007; La Porta et al., 2015; Maldonado et al., 2016). Our results provide evidence that the effect of the CB2r agonist is sufficient to promote drug-taking behavior in nerve-injured mice for alleviation of spontaneous pain, but it is void of reinforcing effects in animals without pain, suggesting the absence of abuse liability. This absence of reinforcement adds value to the modulation of pain through CB2r agonists, since current available agents for neuropathic pain treatment have reduced efficacy and often show addictive properties in humans and rodents (Attal and Bouhassira, 2015; Bonnet and Scherbaum, 2017; Bura et al., 2018; Finnerup et al., 2015; Hipolito et al., 2015; O’Connor et al., 2011).

A previous work using the CB2r agonist AM1241 showed drug-taking behavior and antinociception in nerve-injured rats (Gutierrez et al., 2011), although a recent multicenter study demonstrated off-target effects of this drug (Soethoudt et al., 2017). The disruption of JWH133 effects observed in constitutive knockout mice confirms that the relief of spontaneous pain and the effects reducing mechanical nociception and anxiety-like behavior are mediated by CB2r stimulation. However the CB2r agonist partially preserved its effects promoting drug self-administration and relieving thermal hypersensitivity in CB2KO mice, suggesting that JWH133 may also act through other receptors. JWH133 has shown effects interacting with the Transient Receptor Potential Ankyrin1 (TRPA1) (Soethoudt et al., 2017), a receptor needed for thermal pain perception (Vandewauw et al., 2018), thatcould participate in these responses.

Nerve-injured mice defective in neuronal CB2r showed higher JWH133 intake than wild-type littermates, indicating persistence of drug effects and increased spontaneous pain when neurons do not express CB2r. Importantly, mechanical and thermal neuropathic hypersensitivity before drug self-administration were similar in neuronal knockouts and their wild-type littermates, which suggests different mechanisms of spontaneous pain and evoked nociception. In addition, mechanical nociception measured after JWH133 was more severe in neuronal CB2r knockouts than in wild-type littermates, which indicates decreased JWH133 efficacy on mechanical antinociception. Several studies described the presence of CB2r mRNA and functional CB2r in neuronal populations from different areas of the brain (Stempel et al., 2016; Zhang et al., 2014). However, other works using targeted expression of fluorescent proteins under the control of the mouse gene *Cnr2* failed to describe CB2r expression in neurons (López et al., 2018; Schmöle et al., 2015a). Our results agree with a role of neuronal CB2r during painful neuroinflammatory conditions, a setting that was not studied before in mice defective in neuronal CB2r. Thermal hypersensitivity and anxiety-like behavior measured after self-administration was similar in neuronal knockouts and wild-type mice, which indicates involvement of non-neuronal cell populations. However, it should also be considered that the neuronal knockout mice had higher JWH133 consumption. Thus, a possible lack of efficacy could also be present for thermal antinociception and inhibition of anxiety-like behavior. Although a neuronal involvement was found, CB2r neuronal knockouts did not recapitulate the phenotype of mice constitutively lacking CB2r, suggesting additional cell types involved in the effects of CB2r agonists.

We investigated the effects of JWH133 promoting its own consumption and inducing antinociception and anxiolysis in CB2r LysM Cre+ mice, mainly lacking CB2r in monocytes, the precursors of microglial cells. We did not observe a microglial participation in these pain-related phenotypes, which may be due to an incomplete deletion of CB2r in microglia through LysM-driven Cre expression (Blank and Prinz, 2016). Previous studies in mice constitutively lacking CB2r described an exacerbated spinal cord microgliosis after nerve injury (Nozaki et al., 2018; Racz et al., 2008), which suggested a relevant role of CB2r controlling glial reactivity. Since spinal microgliosis participates in the increased pain sensitivity after a neuropathic insult and macrophages and microglia express CB2r, blunted effects of JWH133 were expected in microglial CB2r knockouts. However, monocyte-derived cells did not seem to be involved in the analgesic effects mediated by the exogenous activation of CB2r in these experimental conditions.

The immunohistochemical analysis of dorsal root ganglia from mice transplanted with bone marrow cells of CB2r GFP BAC mice (Schmöle et al., 2015a) revealed a pronounced infiltration of immune cells expressing CB2r in the dorsal root ganglia after nerve injury. Macrophages and lymphocytes expressing CB2r were found at a time point in which nerve-injured mice present mechanical and thermal hypersensitivity and self-administer compounds with demonstrated analgesic efficacy (Bura et al., 2013, 2018). Interestingly, GFP expression was also found in neurons, suggesting a transfer of CB2r from peripheral immune cells to neurons. An explanation for this finding may come from processes of cellular fusion or transfer of cargo between peripheral blood cells and neurons (Alvarez-Dolado et al., 2003; Ridder et al., 2014). Bone marrow-derived cells fuse with different cell types in a process of cellular repair that increases after tissue damage. These events may be particularly important for the survival of neurons with complex structures that would otherwise be impossible to replace (Giordano-Santini et al., 2016). Alternatively, extracellular vesicles drive intercellular transport between immune cells and neurons (Budnik et al., 2016). Earlier studies showed incidence of fusion events between bone marrow-derived cells and peripheral neurons in a model of diabetic neuropathy (Terashima et al., 2005), and similar processes were observed in central neurons after peripheral inflammation (Giordano-Santini et al., 2016; Ridder et al., 2014). Functional contribution of these mechanisms to neuronal CB2r expression has not yet been explored, although cargo transfer between immune cells and neurons could modify neuronal functionality and it could offer novel therapeutic approaches to modulate neuronal responses (Budnik et al., 2016). Hence, CB2r coming from white blood cells and present in neurons could be significant modulators of spontaneous neuropathic pain and may be contributors to the analgesic effect of CB2r agonists.

Our results suggest participation of lymphoid cells on spontaneous neuropathic pain, but not on basal neuropathic hypersensitivity, highlighting possible differences on the pathophysiology of these nociceptive manifestations. In addition, lymphoid cells were essential for the effects of JWH133 alleviating mechanical sensitivity after a nerve injury. These hypotheses were evaluated by using anti-ICAM1 antibodies that impair lymphocyte extravasation. Previous studies revealed that anti-ICAM1 treatment inhibited opioid-induced antinociception in a model of neuropathic pain (Celik et al., 2016; Labuz et al., 2009). According to the authors, stimulation of opioid receptors from the immune cells infiltrating the injured nerve evoked the release of opioid peptides that attenuated mechanical hypersensitivity. Lymphocyte CB2r could also be involved in the release of leukocyte-derived pain-modulating molecules. Experiments assessing the function of ICAM1 (Celik et al., 2016; Deane et al., 2012; Labuz et al., 2009) showed the participation of this protein on lymphocyte extravasation. In agreement, we observed a decrease of T cell markers in the dorsal root ganglia of mice receiving anti-ICAM1. Anti-ICAM1 treatment also increased the mRNA levels of the B cell marker CD19 in the dorsal root ganglia. Since ICAM1-interacting T cells show activity limiting B cell populations (Deane et al., 2012; Zhao et al., 2006), it is likely that the absence of T cells in the nervous tissue increased infiltration of B lymphocytes. B cells are involved in the severity of neuroinflammatory processes and have been linked to pain hypersensitivity (Huang et al., 2016; Jiang et al., 2016; Li et al., 2014; Zhang et al., 2016). Interestingly, CB2r restrict glucose and energy supply of B cells (Chan et al., 2017), which may alter their cytokine production as previously described for macrophages and T cells. However, the participation of CB2r from B cells on neuropathic pain has not yet been established. Our results indicate an increase in JWH133 consumption that could be driven by an increased infiltration of B cells. Overall, the results with the ICAM-1 experiment suggest a relevant participation of lymphoid CB2r on painful neuroinflammatory responses.

In summary, the contribution of neurons and lymphocytes to the effects of CB2r agonists on spontaneous and evoked pain suggests a coordinated response of both cell types after the nerve injury. CB2r-expressing lymphocytes could participate in pain sensitization through release of pain-related molecules and the observed responses are also compatible with transfer of CB2r between immune cells and neurons. Hence, bone-marrow derived cells may provide a source of functional CB2r that was not considered before and could clarify the controversial presence of these receptors in neurons. Nociceptive and affective manifestations of chronic neuropathic pain are therefore orchestrated through neuronal and immune sites expressing CB2r, highlighting the functional relevance of this cannabinoid receptor in different cell populations.

Our results on operant JWH133 self-administration depict CB2r agonists as candidate painkillers for neuropathic conditions, void of reinforcing effects in the absence of pain. These pain-relieving effects involve the participation of CB2r from neurons and lymphocytes preventing the neuroinflammatory processes leading to neuropathic pain. Therefore, CB2r agonists would be of interest for preventing neuropathic pain development and the potential trials to evaluate this effect should consider starting CB2r agonist treatment before o shortly after the induction of neuropathic insults, as in our study, in contrast to the treatment strategies used in previous clinical trials. The identification of a cannabinoid agonist simultaneously targeting the behavioral traits and the multiple cell types involved in the pathophysiology of chronic neuropathic pain acquires special relevance in a moment in which the absence of efficient painkillers void of abuse liability has become a major burden for public health.

## Materials and Methods

### Animals

C57BL/6J male mice were purchased from Charles River Laboratories (L’Arbresle, France), and knockout male mice were bred in the Institute of Molecular Psychiatry (University of Bonn, Bonn, Germany). CB2r constitutive knockouts were bred from heterozygous parents and their wild-type littermates were used as controls. Neuron and microglia/macrophage-specific conditional CB2r knockout mice were generated as previously described (Stempel et al., 2016). Briefly, mice expressing Cre recombinase under the Synapsin I promoter (Syn) and mice expressing Cre recombinase inserted into the first coding ATG of the lysozyme 2 gene (LysM) were crossed with CB2r floxed animals (Cnr2^fl/fl^ mice). F1 mice were backcrossed to Cnr2^fl/fl^ mice to generate mice Cnr2^fl/fl^ and heterozygous for Cre (Cre-Cnr2^fl/fl^). SynCre-Cnr2^fl/fl^ (Syn Cre+) and LysMCre-Cnr2^fl/fl^ (LysM Cre+) mice were selected and further backcrossed to Cnr2^fl/fl^ mice to produce experimental cohorts containing 50% conditional knockout animals (also referred to as neuronal and microglial knockouts throughout the study) and 50% littermate control animals (referred to as Cre-Negative mice throughout the study). For bone-marrow transplantation studies, 2 CB2-GFP BAC mice (Schmöle et al., 2015a) or C57BL/6J mice were used as donors and C57BL/6J mice were used as recipient mice. All mice had a C57BL/6J genetic background. The behavioral experimental sequence involving operant self-administration and assessment of nociceptive and anxiety-like behavior was repeated 3 times in the experiments assessing the effects of JWH133 doses (Figure 1) and 4 and 5 times in the experiments evaluating constitutive and conditional knockout mice, respectively (Figures 2 and 3). The experiments involving bone-marrow transplantation and lymphocyte depletion were performed once. Sample size was based on previous studies in our laboratory using comparable behavioral approaches (Bura et al., 2013, 2018; La Porta et al., Pain 2015).

The behavioral experiments were conducted in the animal facility at Universitat Pompeu Fabra (UPF)-Barcelona Biomedical Research Park (PRBB; Barcelona, Spain). Mice were housed in a temperature (21±1°C) and humidity-controlled (55±10%) room and handled during the dark phase of a 12h light/dark reverse cycle (light off at 8:00 a.m., light on at 8:00 p.m.). Before starting the experimental procedure, mice were single housed and handled/habituated for 7 days. Food and water were available *ad libitum* except during the training period for food-maintained operant behavior, when mice were exposed to restricted diet for 8 days. Animal handling and experiments were in accordance with protocols approved by the respective Animal Care and Use Committees of the PRBB, Departament de Territori i Habitatge of Generalitat de Catalunya and the Institute of Molecular Psychiatry and were performed in accordance with the European Communities Council Directive (2010/63/EU). Whenever possible, animals were randomly assigned to their experimental condition, and experiments were performed under blinded conditions for surgery and pharmacological treatment (Figure 1), genotype (Figures 2 and 3), bone-marrow transplant and surgery (Figure 4), and antibody treatments (Figure 5).

### Drugs

JWH133 (Tocris, Bristol, UK) was dissolved in vehicle solution containing 5% dimethyl sulfoxide (Scharlab, Sentmenat, Spain) and 5% cremophor EL (Sigma-Aldrich, Steinheim, Germany) in sterilized water and filtered with a 0.22 µm filter (Millex GP, Millipore, Cork, Ireland). JWH133 was self-administered intravenously (i.v.) at 0.15 or 0.3 mg/kg/infusion in volume of 23.5 μl per injection. Thiopental (Braun Medical, Barcelona, Spain) was dissolved in saline and administered through the implanted i.v. catheter at 10 mg/kg in a volume of 50 μl.

### Antibody treatment

Anti-ICAM-1 antibody (clone 3E2; 150 μg; BD Biosciences, Franklin Lakes, NJ, USA) and control rabbit IgG (150 μg; Sigma-Aldrich) were dissolved in saline up to a volume of 300 µl as previously reported (Labuz et al., 2009), and administered i.p. once a day from the day of the surgery to the last self-administration day.

### Operant self-administration

Mice were first trained for operant food self-administration to facilitate subsequent drug self-administration, as previously described (Bura et al., 2018). Briefly, mice were food-restricted for 3 days to reach 90% of their initial weight. Then, mice were trained in skinner boxes (model ENV-307A-CT, Med Associates Inc., Georgia, VT, USA) for 5 days (1 h session per day) to acquire an operant behavior to obtain food pellets (Figure 1–figure supplement 1B, Figure 2–figure supplement 1B, Figure 3– figure supplement 1B, Figure 5–figure supplement 1B). A fixed ratio 1 schedule of reinforcement (FR1) was used, i.e., 1 nose-poke on the active hole resulted in the delivery of 1 reinforcer together with a light-stimulus for 2 s (associated cue). Nose poking on the inactive hole had no consequence. Each session started with a priming delivery of 1 reinforcer and a timeout period of 10 s right after, where no cues and no reward were provided following active nose-pokes. Food sessions lasted 1 h or until mice nose-poked 100 times on the active hole, whichever happened first. After the food training, mice underwent a partial sciatic nerve ligation (PSNL) or a sham surgery, and 4 days later an i.v. catheter was implanted in the right jugular vein to allow drug delivery. Mice started the drug self-administration sessions 7 days after the PSNL/sham surgery. In these sessions, the food reinforcer was substituted by drug/vehicle infusions. Self-administration sessions were conducted during 12 consecutive days, and mice received JWH133 (0.15 or 0.3 mg/kg) or vehicle under FR1 (Figure 1–figure supplement 1B, Figure 2–figure supplement 1B, Figure 3– figure supplement 1B, Figure 5–figure supplement 1B). Sessions lasted 1 h or until 60 active nose-pokes. Active and inactive nose-pokes were recorded after each session and discrimination indices were calculated as the difference between the nose pokes on the active and the inactive holes, divided by the total nose pokes. Data from the last 3 drug self-administration sessions was used for statistical analysis to exclude interference with food-driven operant behavior.

### Partial Sciatic Nerve Ligation

Mice underwent a partial ligation of the sciatic nerve at mid-thigh level to induce neuropathic pain, as previously described (Malmberg and Basbaum, 1998) with minor modifications. Briefly, mice were anaesthetized with isoflurane (induction, 5% V/V; surgery, 2% V/V) in oxygen and the sciatic nerve was exposed at the level of the mid-thigh of the right hind leg. At ∼1 cm proximally to the nerve trifurcation, a tight ligature was created around 33–50% of the cranial side of the sciatic nerve using a 9–0 non-absorbable virgin silk suture (Alcon Cusí SA, Barcelona, Spain) and leaving the rest of the nerve untouched. The muscle was then stitched with 6-0 silk (Alcon Cusí), and the incision was closed with wound clips. Sham-operated mice underwent the same surgical procedure except that the sciatic nerve was not ligated.

### Catheterization

Mice were implanted with indwelling i.v. silastic catheter, as previously reported (Soria et al., 2005). Briefly, a 5.5 cm length of silastic tubing (0.3 mm inner diameter, 0.64 mm outer diameter; Silastic®, Dow Corning Europe, Seneffe, Belgium) was fitted to a 22-gauge steel cannula (Semat Technical Ltd., Herts, UK) that was bent at a right angle and then embedded in a cement disk (Dentalon Plus, Heraeus Kulzer, Wehrheim, Germany) with an underlying nylon mesh. The catheter tubing was inserted 1.3 cm into the right jugular vein and anchored with suture. The remaining tubing ran subcutaneously to the cannula, which exited at the midscapular region. All incisions were sutured and coated with antibiotic ointment (Bactroban, GlaxoSmithKline, Madrid, Spain).

### Nociception

Sensitivity to heat and mechanical stimuli were used as nociceptive measures of neuropathic pain. Ipsilateral and contralateral hind paw withdrawal thresholds were evaluated the day before, 3 and 6 days after the nerve injury, as well as after the last self-medication session on day 18. Heat sensitivity was assessed by recording the hind paw withdrawal latency in response to radiant heat applied with the plantar test apparatus (Ugo Basile, Varese, Italy) as previously reported (Hargreaves et al., 1988). Punctate mechanical sensitivity was quantified by measuring the withdrawal response to von Frey filament stimulation through the up–down paradigm, as previously reported (Chaplan et al., 1994). Filaments equivalent to 0.04, 0.07, 0.16, 0.4, 0.6, 1 and 2 g were used, applying first the 0.4 g filament and increasing or decreasing the strength according to the response. The filaments were bent and held for 4-5 s against the surface of the hindpaws. Clear paw withdrawal, shaking or licking was considered a nociceptive-like response.

### Anxiety-like behavior

Anxiety-like behavior was evaluated with an elevated plus maze made of Plexiglas and consisting of 4 arms (29 cm long x 5 cm wide), 2 open and 2 closed, set in cross from a neutral central square (5 x 5 cm) elevated 40 cm above the floor. Light intensity in the open and closed arms was 45 and 5 lux, respectively. Mice were placed in the neutral central square facing 1 of the open arms and tested for 5 min. The percentage of entries and time spent in the open and closed arms was determined.

### RNA extraction and reverse transcription

Ipsilateral L3-L4 dorsal root ganglia from mice of the ICAM-1 experiment were collected on day 20 after the PSNL. Samples were rapidly frozen in dry ice and stored at −80°C. Isolation of total RNA was performed using the RNeasy Micro kit (Qiagen, Stokach, Germany) according to the manufacturer’s instructions. Total RNA concentration was measured using a NanoDrop ND-1000 Spectrophotometer (NanoDrop Technologies Inc., Montchanin, DE, USA). RNA quality was determined by chip-based capillary electrophoresis using an Agilent Bioanalyzer 2100 (Agilent, Palo Alto, CA, USA). Reverse transcription was performed using Omniscript reverse transcriptase (Qiagen) at 37°C for 60 min.

### Quantitative real-time PCR analysis

The qRT-PCR reactions were performed using Assay-On-Demand TaqMan probes: Hprt1 Mm01545399_m1, CD2 Mm00488928 m1, CD4 Mm00442754_m1, CD19 Mm00515420_m1, C1q Mm00432162_m1 (Applied Biosystems, Carlsbad, CA, USA) and were run on the CFX96 Touch Real-Time PCR machine (BioRad, Hercules, CA, USA). Each template was generated from individual animals, and amplification efficiency for each assay was determined by running a standard dilution curve. The expression of the Hprt1 transcript was quantified at a stable level between the experimental groups to control for variations in cDNA amounts. The cycle threshold values were calculated automatically by the CFX MANAGER v.2.1 software with default parameters. RNA abundance was calculated as 2−(Ct). Levels of the target genes were normalized against the housekeeping gene, Hprt1, and compared using the ΔΔCt method (Livak and Schmittgen, 2001).

### Bone marrow transplantation

C57BL/6J mice received bone marrow from CB2-GFP BAC or C57BL/6J male mice. G-irradiation of C57BL/6J recipient male mice (9.5 Gy) was performed in a 137Cs-g IBL 437C H irradiator (Schering CIS Bio international) at 2.56 Gy/min rate in order to suppress their immune response. Afterwards, approximately 5×10^5^ bone marrow cells collected from donors (CB2-GFP BAC or C57BL/6J) and transplanted through the retro-orbital venous sinus of the recipient mice. Irradiated mice were inspected daily and were given 150 ml of water with enrofloxacin at 570 mg/l and pH 7.4 (Bayer, Germany) for 30 days to reduce the probability of infection from opportunistic pathogens. Peripheral blood samples (150 μl) were collected by tail bleeding into a tube with 0.5 M EDTA solution to evaluate immune system recovery through flow cytometry 4, 8 and 12 weeks after the bone marrow transplantation.

### Flow cytometry

For the analyses of hematopoietic cells, a hypotonic lysis was performed to remove erythrocytes. 50μl of blood was lysed using 500μl of ACK (Ammonium-Chloride-Potassium) Lysing Buffer (Lonza, Walkersville, USA) 10 min at room temperature. After the erythrocytes lysis, 2 washes with PBS were performed prior the incubation with the antibodies for 30 min at 4°C. Cells were stained with the following fluorochrome-coupled antibodies: Allophycocyanin (APC)-conjugated anti-mouse CD11b (1:300; cn.17-0112 eBioscience, USA) to label myeloid cells, phycoerythrin (PE)-conjugated anti-mouse B220 (1:100; cn.12-0452, eBioscience, USA) for B lymphocytes and phycoerythrin/cyanine (PE/Cy7)-conjugated anti-mouse CD3, 1:100; cn.100320, BioLegend, USA) for T lymphocytes. Immunofluorescence of labeled cells was measured using a BD™ LSR II flow cytometer. Dead cells and debris were excluded by measurements of forward-versus side-scattered light and DAPI (4′,6-diamino-2-phenylindole) (Sigma) staining. Gates for the respective antibodies used were established with isotype controls and positive cell subset controls. Data analysis was carried out using FACSDiva version 6.2 software (BD biosciences).

### Immunohistochemistry

Mice were sacrificed 2 weeks after the PSNL/sham surgery and L3-L5 dorsal root ganglia were collected to quantify GFP+ cells in mice transplanted with bone marrow cells of CB2-GFP or C57BL6/J mice. Ganglia were freshly extracted and fixed in 4% paraformaldehyde during 25 min at 4°C. After 3×5 min washes with phosphate buffered saline (PBS) 0.1 M (pH 7.4), were preserved overnight in a 30% sucrose solution in PBS 0.1 M containing sodium azide 0.02%. 24 h later, ganglia were embedded in molds filled with optimal cutting temperature compound (Sakura Finetek Europe B.V., Netherlands) and frozen at −80°C. Samples were sectioned with a cryostat at 10 µm, thaw-mounted on gelatinized slides and stored at −20°C until use. Dorsal root ganglia sections were treated 1 h with 0.3 M glycine, 1 h with oxygenated water 3% (Tyramide Superboost Kit, B40922, Thermo Fisher, USA) and, after 3×5 min washes with PBS 0.01 M, 1 h with blocking buffer. Samples were incubated 16 h at room temperature with rabbit anti-GFP (1:2000, A11122, Thermo Fisher, USA) and chicken anti-neurofilament heavy (NFH) (1:1000, ab4680, Abcam, UK) antibodies. After 3×10 min washes with PBS 0.01 M, sections were incubated with anti-rabbit poly-HRP-conjugated secondary antibody for 1 h and washed 4×10 min. Alexa Fluor™ tyramide reagent was applied for 10 min and then the Stop Reagent solution for 5 min (Tyramide Superboost Kit). Afterwards samples were incubated 2 h at room temperature with primary antibodies diluted in blocking buffer (PBS 0.01 M, Triton X-100 0.3%, Normal Goat Serum 10%). The following primary antibodies were used: rabbit anti-peripherin (1:200, PA3-16723, Thermo Fisher, USA), rabbit anti-β-III tubulin (1:1000, ab18207, Abcam, UK), rat anti-CD45R/B220 APC (1:500, Clone RA3-6B2, 103229, Biolegend, USA) and rat anti-F4/80 (1:500, Clone A3-1, MCA497GA, Biorad, USA). After 3×5 min washes, all sections were treated with goat secondary antibodies from Abcam (UK) for 1 h at room temperature: anti-chicken Alexa Fluor® 647 (1:1000, ab150171), anti-rabbit Alexa Fluor® 555 (1:1000, ab150078) and anti-rat Alexa Fluor® 555 (1:1000, ab150158). Samples were then washed with PBS 0.01 M and mounted with 24×24 mm coverslips (Brand, Germany) using Fluoromount-G with DAPI (SouthernBiotech, USA).

### Microscope image acquisition and processing

Confocal images were taken with a Leica TCS SP5 confocal microscope (Leica Microsystems, Mannheim, Germany) on a DM6000 stand using 20x 0.7 NA Air and 63x 1.4 NA Oil Immersion Plan Apochromatic lenses. Leica Application Suite Advanced Fluorescence software (Leica Microsystems, Mannheim, Germany) was used to acquire the images and DAPI, Alexa 488 and Alexa 555 channels were taken sequentially. Images of DAPI were taken with 405 nm excitation and emission detection between 415 and 480 nm; images of Alexa 488 were taken with 488 nm excitation and emission detection between 495 and 540 nm; and images of Alexa 555 were taken with 543 nm excitation and emission detection between 555 and 710 nm. Room temperature was kept at 22±1°C during all imaging sessions. All images were equally processed and quantified with Fiji software (National Institutes of Health, USA). To determine the percentage of dorsal root ganglia area occupied by GFP (+) neurons auto-threshold (“Otsu”) was set between 0-50 in all images and then converted to mask. Afterwards, operations included Close, Fill holes and Watershed neurons for separation and particles between 100-100000 pixel units and circularity 0.2-1.0 were counted. To analyze the number of GFP+ cells per dorsal root ganglia area, background was subtracted from all images (rolling=5), set to an auto-threshold (“Default”) between 0-70 and converted to mask. Particles considered GFP+ were 7-100 microns2 and 0.2-1.0 circularity.

### Statistical analysis

Self-administration and nociceptive behavioral data were analyzed using a linear mixed model with 3 (surgery, day and dose) or 2 factors (day and genotype or antibody treatment) and their interactions. For the covariance structure of the repeated measures, a diagonal matrix was chosen. Bonferroni post hoc analysis was performed when pertinent. Areas Under the Curve (AUCs) of time-courses for operant responding were analyzed using 2-way analysis of variance (ANOVA). Active and inactive responses were analyzed taking into account surgery and dose effects in the dose-response experiments, and active/inactive and genotype or antibody treatment in the knockout and antibody experiments. Anxiety-like behavior was analyzed using 2-way ANOVA (surgery and dose for dose-response experiments), 1-way ANOVA (genotype of conditional knockouts) or t-tests (constitutive knockout and antibody treatment), followed by Bonferroni adjustments when required. Immunohistochemistry of bone marrow-transplanted mice was analyzed using Kruskal Wallis non-parametric tests followed by Mann–Whitney U tests (non-gaussian distribution revealed by Kolmogorov-Smirnov normality test), and qPCR results after antibody treatments were compared with t-tests. IBM SPSS 19 (SPSS Inc., Chicago, IL, USA) and STATISTICA 6.0 (StatSoft, USA) software were used to analyze the data, and differences were considered statistically significant when p value was below 0.05. All experimental data and statistical analyses of this study are included in the manuscript and its supplementary files. Raw data and results of statistical analyses are provided in the Source Data File and its containing data sheets.

## Acknowledgements

Financial support of European Commission [NeuroPain, FP7-602891-2], Instituto de Salud Carlos III, Redes temáticas de investigación cooperativa en salud – Red de trastornos adictivos [#RD12/0028/0023/FEDER], “Ministerio de Economía y Competitividad-MINECO” [#SAF2014-59648-P], “Generalitat de Catalunya-Agència de Gestió d’Ajuts Universitaris i de Recerca-AGAUR” [#2014-SGR-1547 and #2018 FI_B 00207] and “AGAUR” [Institució Catalana de Recerca i Estudis Avançats Academia Award 2015] to R.M. and Polish Ministry of Science and Education [#3070/7.PR/2014/2]. Authors thank Itzel M. Lara, Hugo Ramos, Roberto Cabrera and Cristina Fernández for their help and technical expertise.

## Competing Interests

The authors declare no conflict of interest.

## Supplementary figures

**Figure 1-figure supplement 1.**
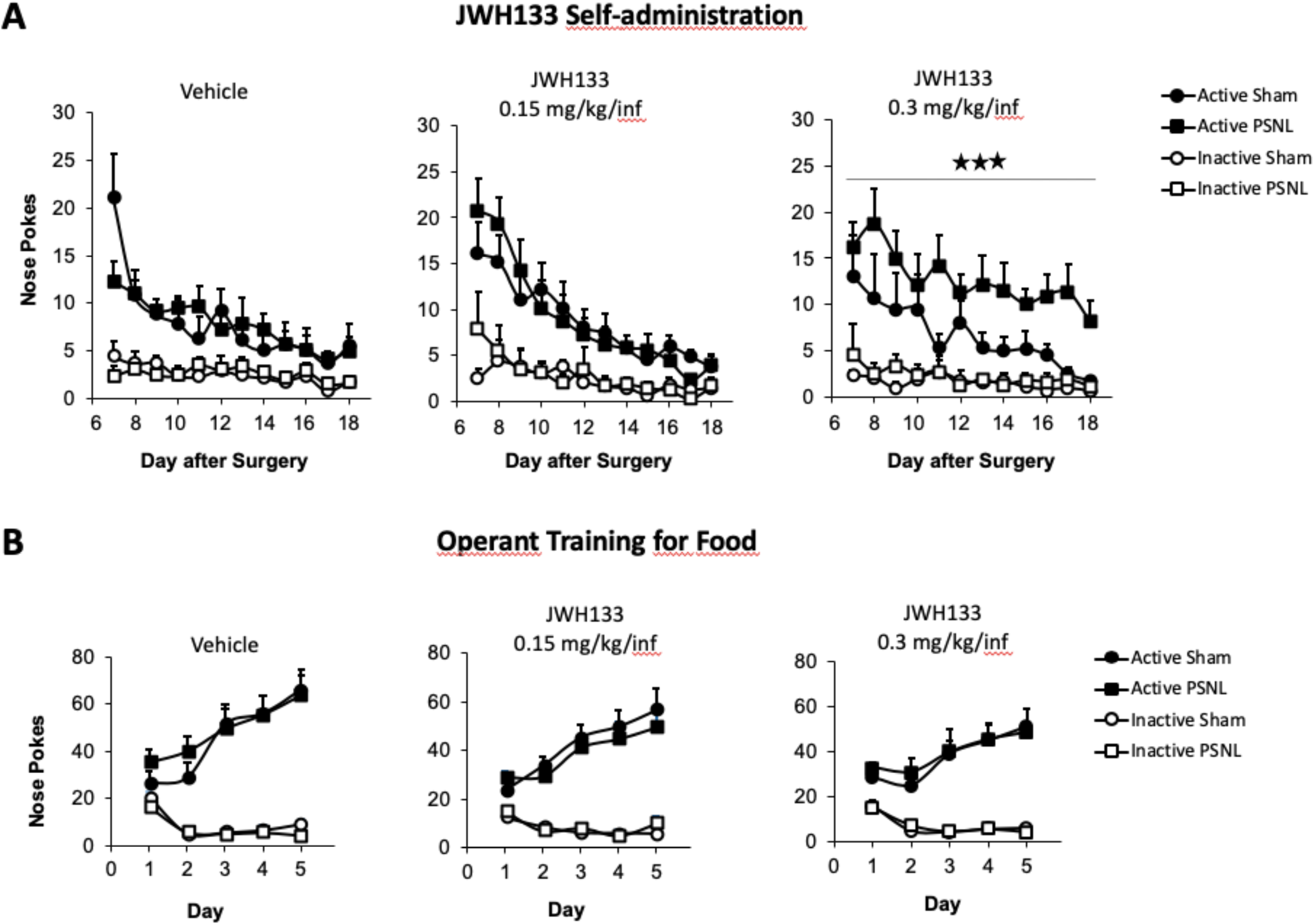
JWH133 self-administration after nerve injury or sham surgery in C57BL6/J mice and food-maintained operant training before the drug self-administration. **A)** The first day of i.v. self-administration, sham-operated mice exposed to the vehicle showed higher active nose pokes than nerve-injured mice exposed to the same treatment. For the rest of the JWH133 self-administration period, mice exposed to the vehicle or to the low dose of JWH133 (0.15 mg/kg/inf) showed similar operant behaviour, regardless of the type of surgery. Nerve-injured mice exposed to the high dose of JWH133 (0.3 mg/kg/inf.) showed higher active nose poking than sham mice exposed to this dose. Inactive responding was similar regardless of type of the surgery and treatment. **B)** All groups of mice developed operant behaviour directed to obtain food pellets before the partial sciatic nerve ligation (PSNL) or the sham surgery. N=7-10 mice per group. Mean and error bars representing SEM are shown. Stars represent p<0.001 vs. respective sham group. Raw data and statistics available in Source Data File.

**Figure 2-figure supplement 1.**
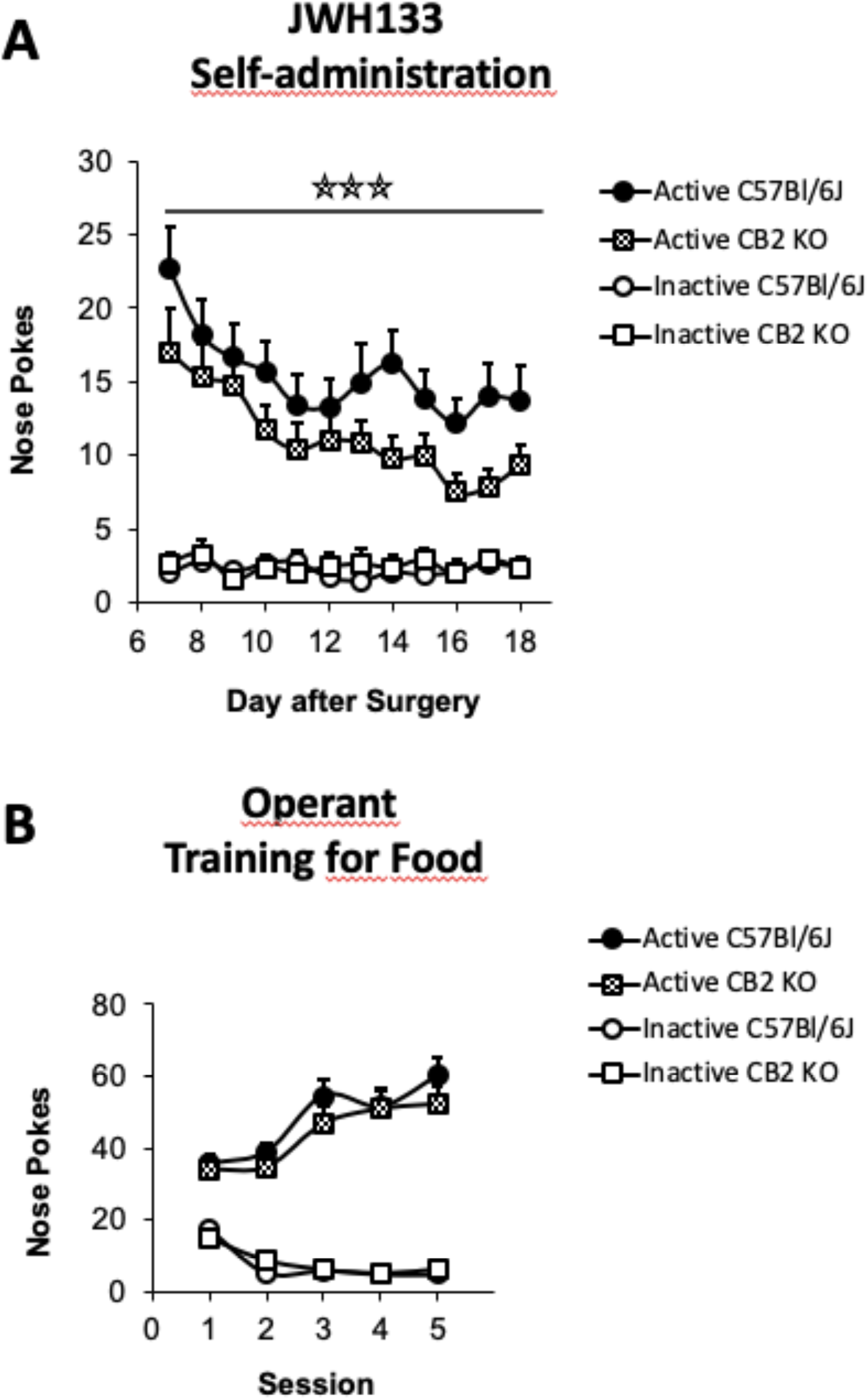
JWH133 self-administration in C57BL6/J and CB2r constitutive knockout (CB2 KO) mice and food-maintained operant training before nerve injury and drug self-administration. **A)** Nerve-injured CB2 KO mice showed a disruption of the active operant behaviour directed to obtain high doses of the CB2r agonist JWH133 (0.3 mg/kg/inf.). **B)** C57BL6/J and CB2 KO mice developed similar operant behaviour for food before the partial sciatic nerve ligation. N=16-19 mice per group. Stars represent p<0.001 vs. C57Bl6/J. Raw data and statistics available in Source Data File

**Figure 3-figure supplement 1.**
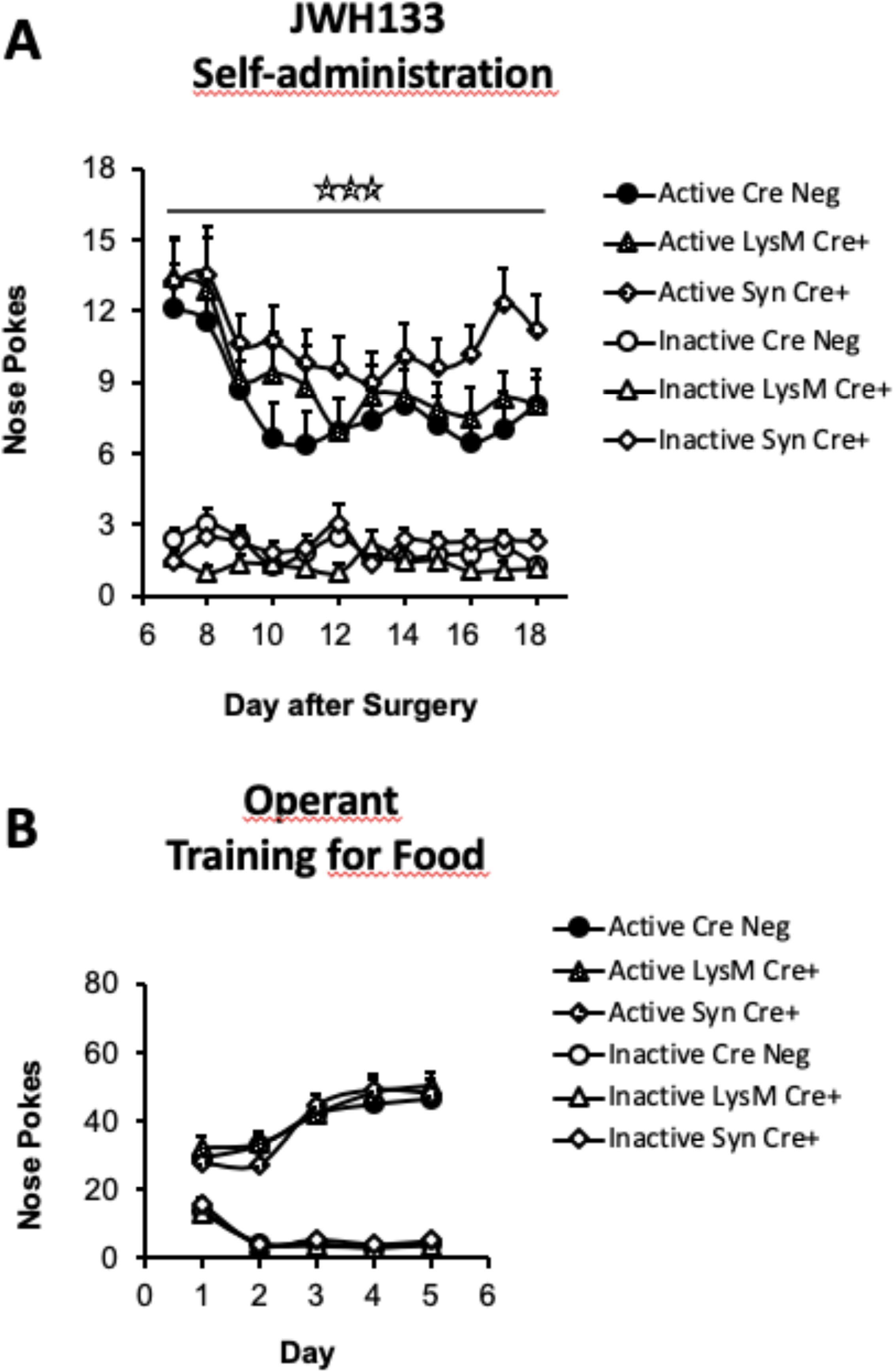
JWH133 self-administration in mice lacking CB2r in neurons or monocytes and their wild-type littermates and food-maintained operant training before nerve injury and drug self-administration. **A)** Mice lacking CB2r in neurons (Syn Cre+) mice showed increased active operant behaviour directed to obtain high doses of the CB2r agonist JWH133 (0.3 mg/kg/inf.). Operant responding for the CB2r agonist was similar between mice lacking CB2r in monocytes (LysM Cre+) mice and their wild-type littermates (Cre Neg). **B)** Syn Cre+, LysM Cre+ and Cre Neg mice developed similar operant behaviour for food before the partial sciatic nerve ligation. N=18-36 mice per group. Stars represent p<0.001 vs. Cre Neg. Raw data and statistics available in Source Data File.

**Figure 4-figure supplement 1.**
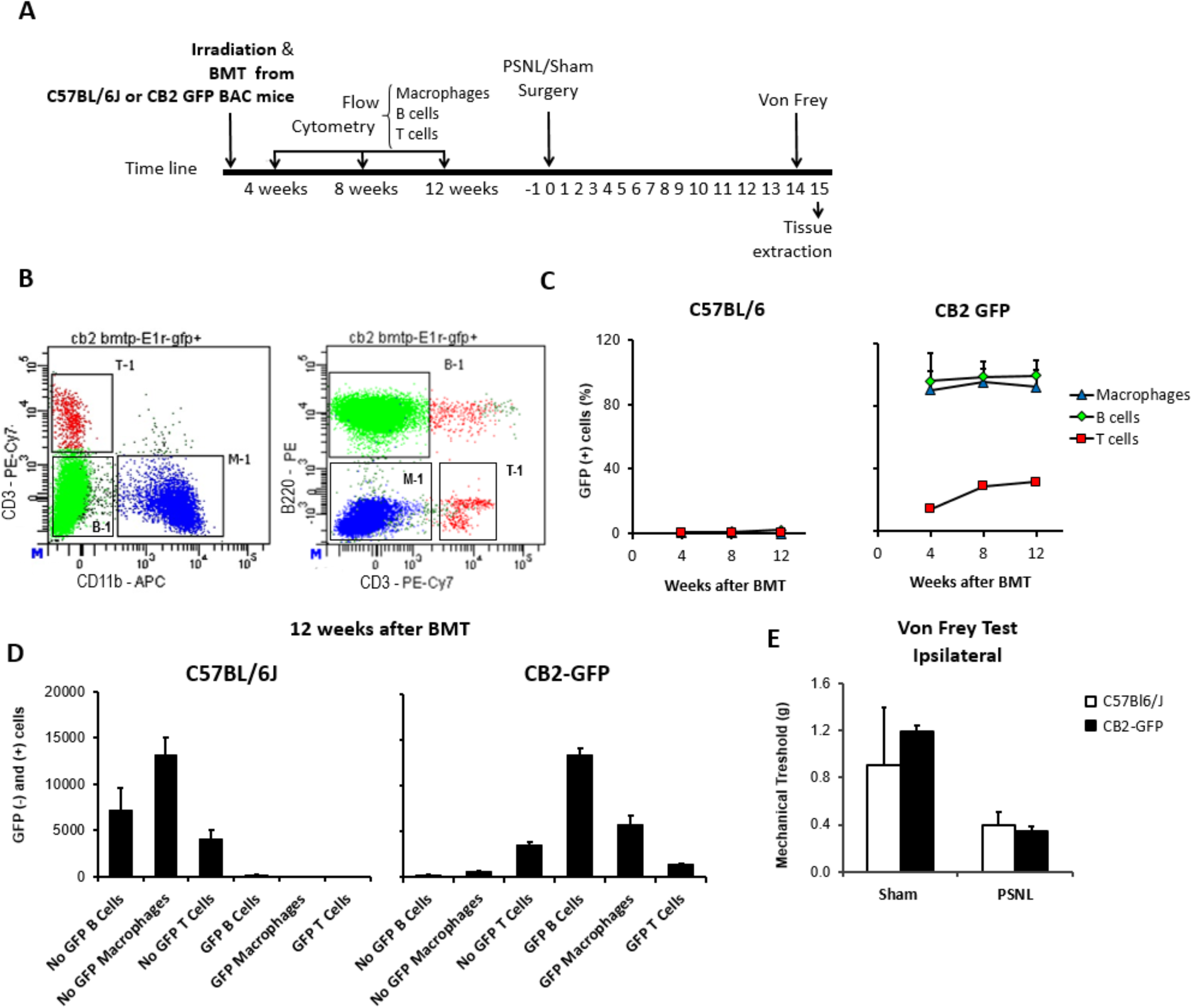
Bone marrow transplantation from CB2 GFP BAC to C57BL6/J mice yields mice with peripheral blood cells expressing GFP. **A)** C57BL6/J mice were irradiated and immediately transplanted with bone marrow cells from CB2 GFP BAC or C57BL6/J mice, yielding CB2-GFP or C57BL6/J mice. Repeated flow cytometry assays were conducted 4, 8 and 12 weeks after the bone marrow transplantation to assess reconstitution of the immune system. Afterwards, a nerve injury or sham surgery was conducted in mice with successful reconstitution (day 0). 14 days later, mechanical nociception was assessed with the von Frey test and dorsal root ganglia samples were collected the following day. **B)** Representative dot plot of flow cytometry showing the labeling of peripheral blood cells from a CB2-GFP bone marrow-transplanted mouse. Cells were pre-gated as single live cells using DAPI staining. T cells (T-1), B cells (B-1) and macrophages (M-1) were gated. PE/Cy7 labeled CD3+ T lymphocytes, APC CD11b+ myeloid cells and PE-B220 labeled B lymphocytes. **C)** Percentage of GFP+ immune cells in C57Bl/6J and CB2-GFP mice from 4 to 12 weeks after transplantation. **D)** 12 weeks after transplantation CB2-GFP mice showed 98% of macrophages GFP+, 86% of B cells GFP+ and 30% of T cells GFP+, whereas C57BL/6J mice did not show significant GFP signal in the different cell populations. n= 4-6 mice per group. **E)** Mechanical thresholds measured before sample collection showed ipsilateral paw sensitization in C57BL/6J and CB2-GFP mice with the nerve injury. n= 2-3 mice per group. Means and error bars representing SEM are shown. Raw data and statistics available in Source Data File.

**Figure 4-figure supplement 2.**
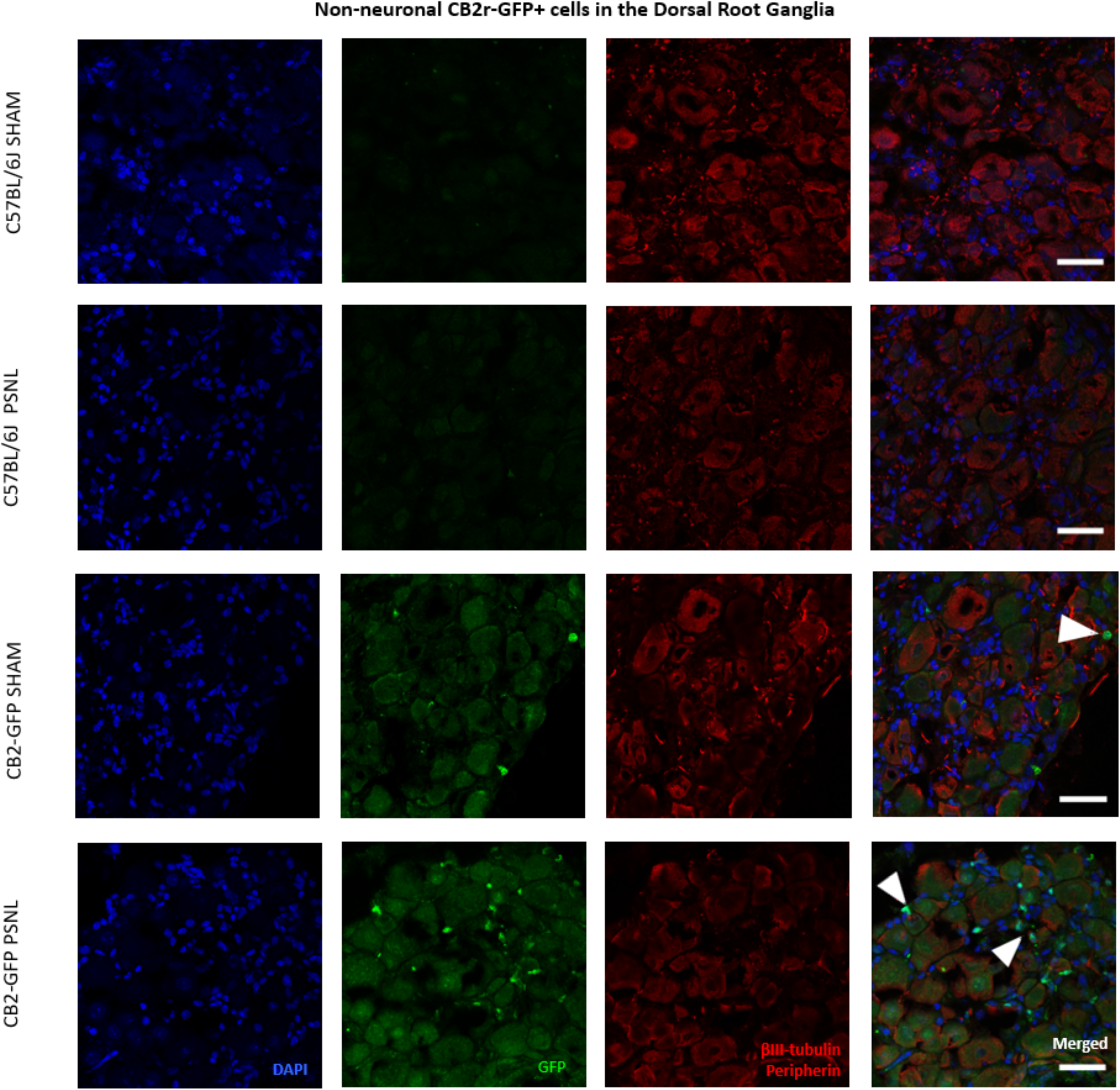
Non-neuronal CB2r-GFP+ cells in the Dorsal Root Ganglia. Split and merged channels of L3-L5 dorsal root ganglia images from sham or nerve-injured mice transplanted with bone marrow cells from CB2 GFP BAC mice (CB2-GFP) or C57BL6/J mice (C57BL6/J). Dorsal root ganglia sections were stained with the nuclear marker DAPI, anti-GFP, and neuronal markers anti-β-III tubulin and anti-peripherin. Sham or nerve-injured C57BL6/J mice did not show significant GFP immunorreactivity. CB2-GFP mice showed infiltration of bone-marrow derived cells enhanced with the nerve injury. Scale bar, 45 μm. White arrows point to GFP+ cells. A certain degree of image processing has been applied equally across merged images to allow optimal visualization.

**Figure 4-figure supplement 3.**
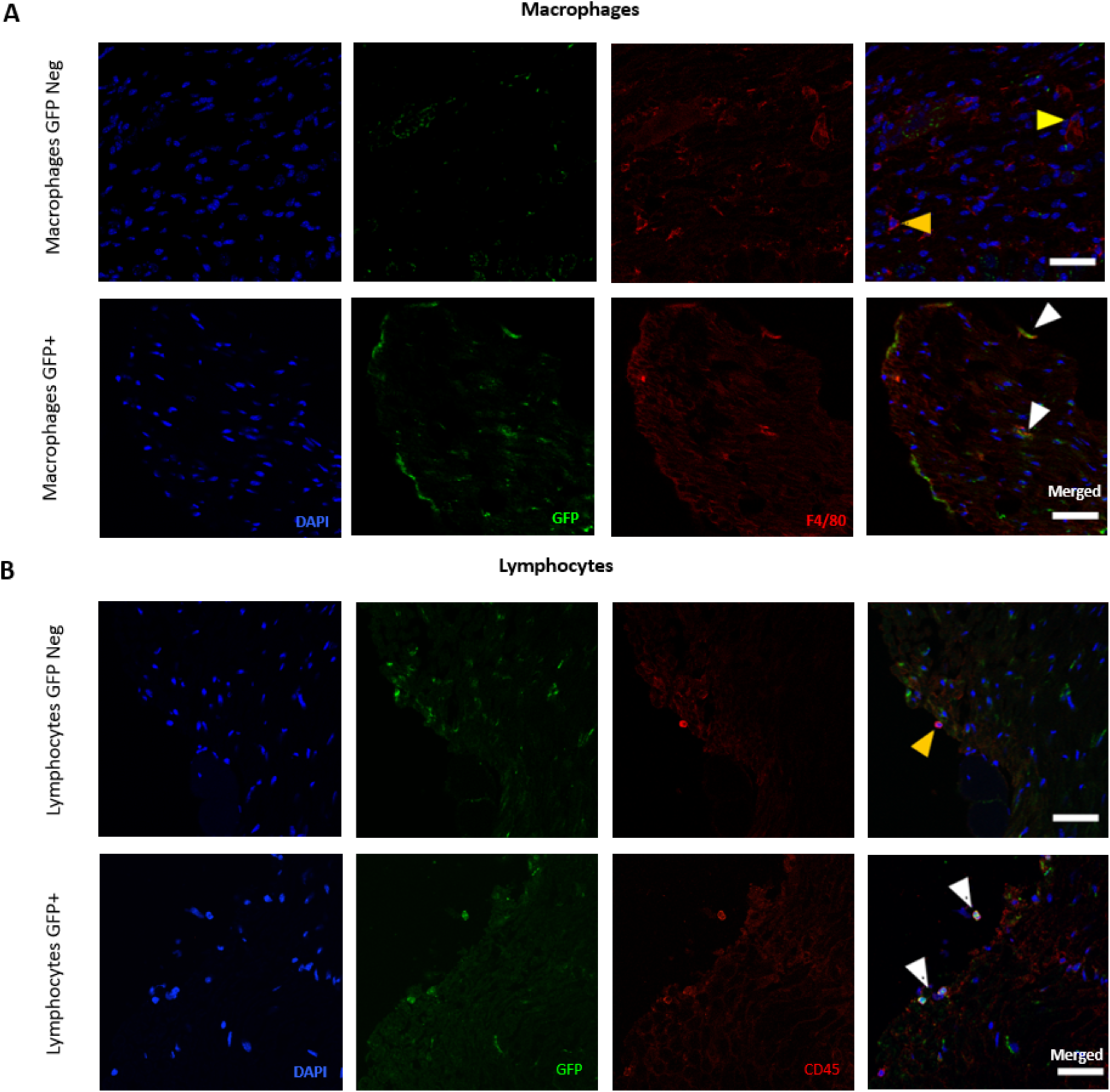
Presence of CB2r-GFP in immune cells in the Dorsal Root Ganglia. Split and merged channels showing **A)** Co-localization of CB2-GFP and the macrophage marker anti-F4/80. **B)** Co-staining of anti-GFP and lymphocyte marker anti-CD45. Scale bar, 45 μm. Yellow arrows point to GFP negative cells and white arrows to GFP+ cells. Certain degree of image processing has been applied equally across merged images for optimal visualization.

**Figure 4-figure supplement 4.**
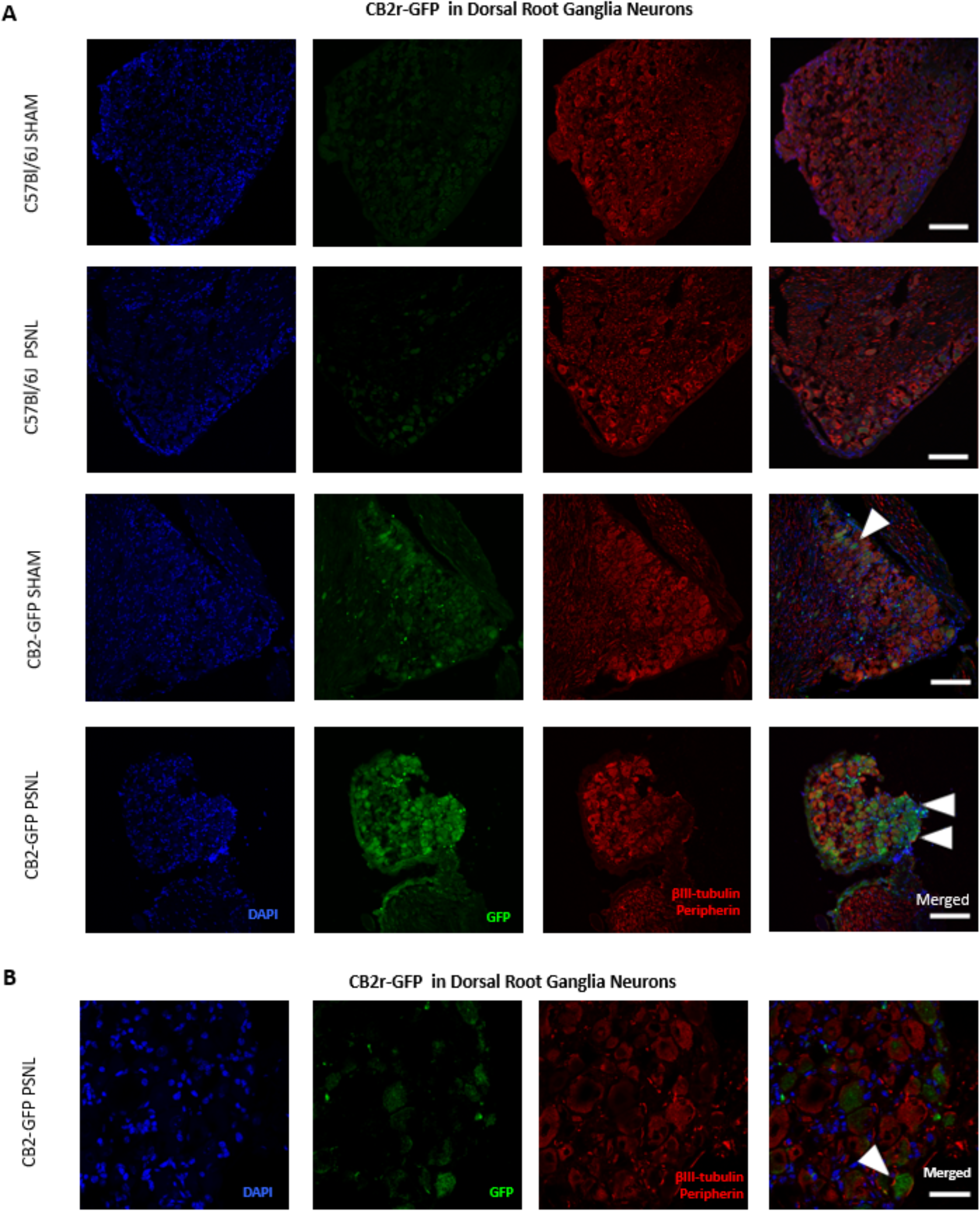
CB2r-GFP in Dorsal Root Ganglia Neurons. Images of L3-L5 dorsal root ganglia from sham or nerve-injured mice transplanted with bone marrow cells from CB2 GFP BAC mice (CB2-GFP) or C57BL6/J mice (C57BL6/J). **A)**, **B)** Split and merged channels of dorsal root ganglia sections stained with the nuclear marker DAPI, anti-GFP, and neuronal markers anti-β-III tubulin and anti-peripherin. **A)** Sham or nerve-injured C57BL6/J mice did not show significant GFP immunorreactivity. CB2-GFP mice showed a percentage of GFP+ neurons that was enhanced with the nerve injury. Scale bar, 140 μm. **B)** Amplified section of dorsal root ganglia from CB2-GFP PSNL mice showing neuronal GFP. Scale bar, 45 μm. Certain degree of image processing has been applied equally across the merged images for optimal visualization.

**Figure 5-figure supplement 1.**
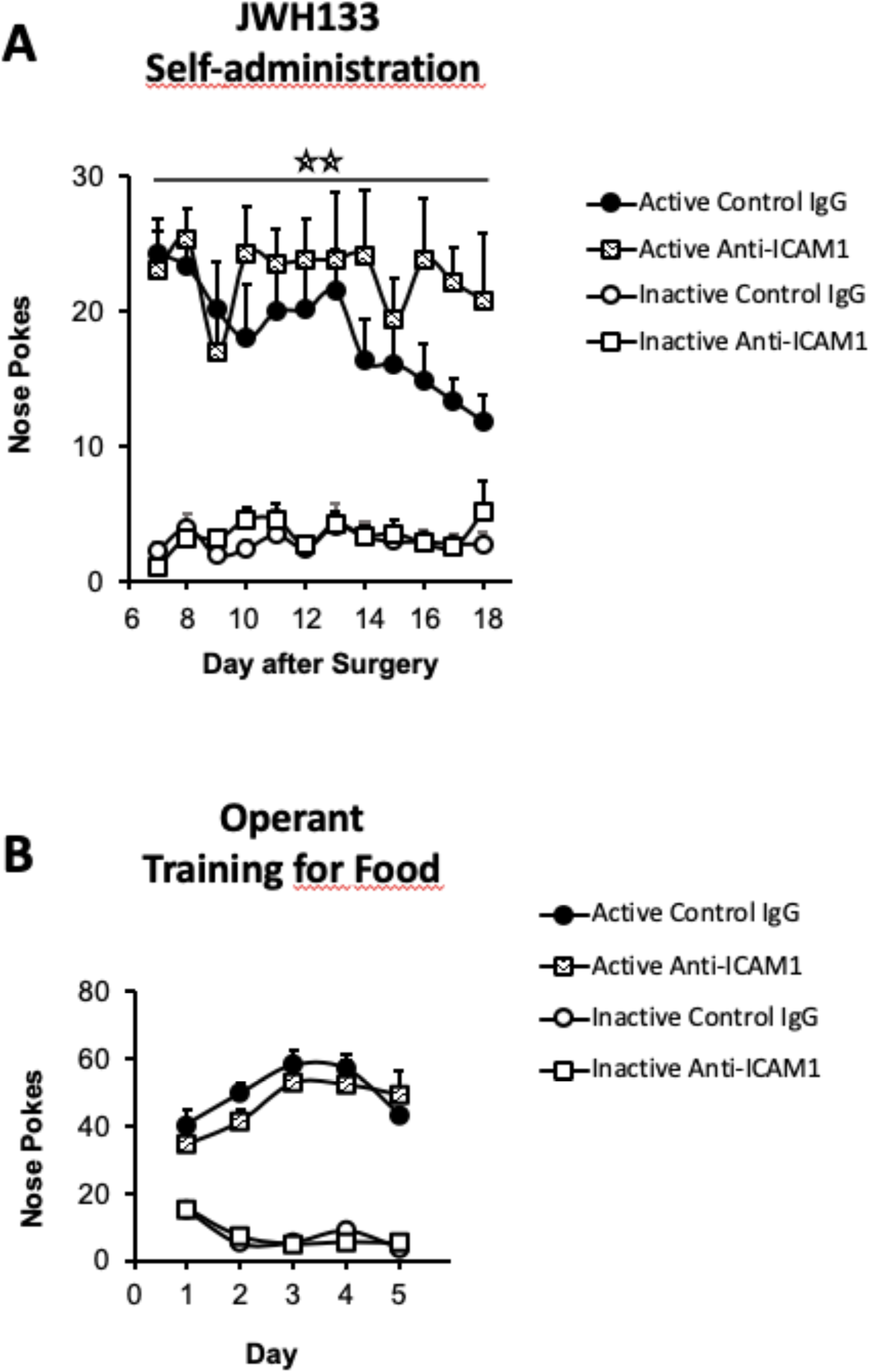
JWH133 self-administration of nerve-injured mice treated with anti-ICAM1 or control IgG and food-maintained operant training before nerve injury and drug self-administration. A) Anti-ICAM1-treated mice showed increased active operant responding directed to obtain high doses of the CB2r agonist JWH133 (0.3 mg/kg/inf.) after the nerve injury. B) C57BL6/J mice of both groups anti-ICAM1 and control IgG developed similar operant behaviour for food before the partial sciatic nerve ligation. N=6-7 mice per group. Stars represent p<0.01 vs. control IgG group. Raw data and statistics available in Source Data File.

